# Tissue-Specific Regulatory and Expression Patterns of CDG-causative genes account to Phenotypic Variability

**DOI:** 10.1101/2025.10.09.681373

**Authors:** Cátia José Neves, António Gomes, Rita Adubeiro Lourenço, Ana Rita Grosso, Paula Videira

## Abstract

Congenital disorders of glycosylation (CDGs) are rare metabolic diseases caused by impaired addition of glycans to proteins and lipids. They display wide clinical variability, but the molecular basis remains unclear. Using healthy transcriptomic data from the GTEx project (bulk: 2,833 samples across 36 tissues; single-nucleus: 209,126 nuclei from 25 samples spanning 8 tissue types and 44 cell types), we investigated the expression and regulatory landscapes of 12 CDG-causative genes.

CDG-causative genes were broadly but heterogeneously expressed, with both high and low expression aligning with clinical features. Interindividual variability in expression may contribute to phenotypic diversity in CDG. *VPS13B* was broadly expressed in fibroblasts, supporting their use in patient-derived models, while other genes showed more tissue-restricted expression, underscoring the importance of cellular context.

Several genes exhibited tissue-specific deviations from balanced biallelic expression. Five allelic expression types across individuals - biallelic balanced or biased, tissue-specific or constitutive monoallelic, and autosomal random monoallelic - suggest a role for allelic regulation in phenotypic heterogeneity.

Tissue-specific eQTLs highlighted regulatory complexity, with many variants located in intronic enhancers of unrelated genes.

Finally, gene expression-immune cell correlations recapitulated known immune phenotypes and suggested context-dependent immune roles.

Together, these findings reveal how genetic, regulatory, and immune factors shape CDG heterogeneity and provide a framework for future research.

## Introduction

Glycosylation is a fundamental biological process in which sugar chains (glycans) are covalently attached to proteins and lipids, altering their structure, solubility, stability, function, and subcellular localization (Eichler, 2019; Varki & Kornfeld, 2022). These modifications are essential for a wide range of physiological processes such as cell-cell and cell-matrix interactions, intracellular signalling, immune recognition, and pathogen detection (Pinho et al., 2023).

Congenital disorders of glycosylation (CDGs) are a group of rare, inherited metabolic diseases caused by genetic defects in enzymes involved in glycosylation pathways (Ng et al., 2024). To date, around 200 CDG types have been described (Ng et al., 2024), typically named after the affected gene (e.g., PMM2-CDG) (Jaeken et al., 2009). These disorders show remarkable clinical variability, with onset ranging from the prenatal period, sometimes resulting in miscarriage, to adulthood (Altassan et al., 2019; Hinderlich et al., 2015). While most CDGs present as multisystem disorders, some are limited to a single organ and show a more localized clinical impact (Francisco et al., 2020; Francisco, Marques-da-Silva, et al., 2019; Francisco, Pascoal, et al., 2019). CDGs can result from different types of genetic mutations, sometimes even within the same gene, leading to diverse inheritance patterns and/or clinical outcomes (Francisco et al., 2023). For example, mutations in different regions of the *GNE* gene can cause two distinct forms of GNE-CDG: an autosomal recessive (ar) form known as GNE myopathy, which primarily affects skeletal muscle, and an autosomal dominant (ad) form called Sialuria, which involves multiple organ systems (Hinderlich et al., 2015). Moreover, even individuals carrying the same pathogenic variant may present with variable symptoms and disease progression, as observed in GNE-CDG (ar) (Boyden et al., 2011).

CDG phenotypic variability is not limited to organ-specific manifestations but also extends to immune system involvement, which is increasingly recognized as a key feature in many CDG subtypes. To date, 23 CDG types have been associated with immune-related features, including recurrent infections, inflammation, allergies, autoimmune features, biochemical abnormalities, and unusual responses to vaccination. Of these, 11 types have minor immune involvement (affecting <50% of patients, e.g., ALG1-, MAN1B1- and PMM2-CDG), while 12 show major involvement (affecting >50% of patients, e.g., MOGS-, PGM3- and VPS13B-CDG) and categorized as inborn errors of immunity according to the International Union of Immunological Societies (IUIS) (Monticelli et al., 2016; Pascoal et al., 2020, 2024).

Despite growing knowledge of the genetic causes and clinical features of CDGs, the molecular basis for their tissue-specific manifestations and phenotypic variability remains poorly understood. Such difficulties are compounded by challenges in obtaining biopsies from affected tissues. To address this gap, we systematically analyzed transcriptome data from healthy human tissues (GTEx) to explore gene expression levels, allelic expression patterns, expression quantitative trait loci (eQTLs) and immune-association of CDG-causative genes. To ensure a representative and clinically relevant analysis, we selected 12 CDG types based on their prevalence, organ-specific manifestations, and known immune involvement. This cohort comprises (i) the most prevalent forms with global distribution and multi-organ involvement, including PMM2-, ALG6-, ALG1-, SLC35A2-, ALG13-, SRD5A3-, MAN1B1-, and DPAGT1-CDG (Piedade et al., 2022); (ii) a primarily muscle-specific form, GNE-CDG (ar); and (iii) CDG types with major immune involvement, MOGS-, PGM3-, and VPS13B-CDG (Pascoal et al., 2024).

Our findings underscore the biological relevance of CDG-causative genes, which are broadly expressed across healthy tissues. We found marked variability in gene expression, allele-specific expression, and the number of eQTLs across tissues and individual factors that may contribute to the phenotypic variability observed in CDG. Many eQTLs were located within diverse regulatory elements, particularly intronic enhancers of unrelated genes, underscoring the complexity and long-range regulation of these genes. Notably, we also uncovered strong correlations between CDG-causative gene expression and immune cell abundance in specific tissues, many of which align with those clinically impacted in the corresponding CDG types.

## 2. Methods

### 2.1. Selection of CDG Types and Genes

A total of 12 CDGs were selected for analysis (**Sup Table 1**). Eight of these, ALG1-, ALG13-, ALG6-, DPAGT1-, MAN1B1-, PMM2-, SLC35A2-, and SRD5A3-CDG, were chosen based on their higher prevalence, global distribution, and multi-organ involvement (Piedade et al., 2022). To capture tissue-specific disease, GNE-CDG (ar), which primarily affects skeletal muscle was included (Pereira et al., 2025). Additionally, three CDGs with major immune involvement, MOGS-, PGM3-, and VPS13B-CDG (Pascoal et al., 2024), were included. Ten of the selected CDGs are inherited in an autosomal recessive manner, while two are X-linked, either dominant or recessive (e.g., ALG13-CDG) (Cai et al., 2022), or exclusively X-linked dominant (e.g., SLC35A2-CDG) (Miyamoto et al., 2019).

### 2.2. Transcriptomic Data Acquisition and Processing

Transcriptome profiles from 948 post-mortem adult donors, comprising 17,382 samples across 54 tissue types were obtained via the Genotype-Tissue Expression (GTEx) project (release v8) (Aguet et al., 2017; Carithers et al., 2015; Castel et al., 2020; Consortium et al., 2020). The GTEx datasets were enclosed: bulk gene expression data in transcripts per million (TPM) (GTEx_Analysis_2017-06-05_v8_RNASeQCv1.1.9_gene_tpm.gct.gz); haplotype-resolved expression counts from WASP-corrected RNA-seq alignments with phasing maintained across genes per sample (phASER_WASP_GTEx_v8_matrix.gw_phased.txt.gz); expression quantitative trait loci (eQTL) data for CDG-causative genes; and sample annotations (GTEx_Analysis_v8_Annotations_SampleAttributesDS.txt). GTEx expanded subject phenotype data (phs000424.v8.pht002742.v8.p2.c1.GTEx_Subject_Phenotypes.GRU.txt) were accessed through the NCBI database of Genotypes and Phenotypes (dbGaP) under accession phs000424.v8.p2, in compliance with the NIH Genomic Data Sharing Policy. We integrated all datasets except the eQTLs and considered only paired-end samples with at least 60 million reads per sample and prepared with the Illumina TruSeq library construction protocol (nonstrand specific polyA+ selected library). Cell culture samples were excluded. Healthy subjects were selected by filtering samples for “violent and fast deaths” and “no terminal diseases”. After filtering, 2,833 samples from 36 healthy tissues were retained for downstream gene and allelic expression analyses of 12 CDG-causative genes (**Sup Table 2**). eQTL data for CDG-causative genes were used as provided by GTEx without additional filtering.

Additionally, single-nucleus RNA sequencing (snRNA-seq) data (GTEx_8_tissues_snRNAseq_atlas_071421.public_obs.h5ad) obtained from GTEx included 209,126 single-nucleus transcriptomes from 25 archived frozen tissue samples across 8 tissue types (including skin) from 16 post-mortem adult donors, spanning 44 cell types (Eraslan et al., 2022). Cell-specific datasets were generated by retaining entries with at least three samples and excluding features with all-zero values. Filtered CDG-causative gene expression data were converted to sparse matrices using the Matrix (v1.6.5) R package (Bates et al., 2024). Metadata (tissue types) were structured with cell IDs as row identifiers. These CDG-related sparse matrices and metadata were used to construct Seurat objects and were subsequently normalized using the default pipeline provided by the Seurat (v5.1.0) R package (Hao et al., 2024).

### 2.3. CDG Phenotype to Tissues Involvement Mapping

To identify frequent GTEx tissues associated with the studied CDG types, we retrieved phenotype tables for each CDG from The Human Disease Database (MalaCards). We selected phenotypes classified as: higher than 33% in Human Phenotype Ontology (HPO) or higher than 30% in Orphanet. Frequent phenotypes were manually assigned to one of the GTEx tissues based on their HPO definitions and comments and subsequently reviewed by a clinician (**Sup Table 3**).

### 2.4. Allelic Expression of CDG-causative genes

To explore CDG-causative allele expression patterns across tissues, Minor Allele Frequency (MAF) was calculated for each sample using haplotype-resolved expression counts from WASP-corrected RNA-seq alignments and dividing the Minimum Haplotype Count by the Total Haplotype Count (HAPLOTYPE_1_COUNT + HAPLOTYPE_2_COUNT).

To classify CDG-causative genes according to allele expression types in each individual, we applied a three-step approach: (i) calculation of Allele 1 Ratios, (ii) identification of genes with autosomal random monoallelic expression (aRME) or X-chromosome inactivation based on the method by Kravitz et al. (2023), and (iii) classification of the remaining four allele expression types. Allele 1 Ratios were calculated for each sample using haplotype-resolved expression counts from WASP-corrected RNA-seq alignments, by dividing the Haplotype 1 Count by the total haplotype count.

Following Kravitz et al. (2023), for each individual, allele ratio distributions across ≥3 tissues were tested against a binomial (H₀) versus beta-binomial (H₁) model. Binomial distributions are expected for genes with biallelic balanced expression or consistent silencing of one allele across cells or tissues (e.g., biallelic bias, tissue-specific or constitutive monoallelic expression/ imprinting). In contrast, beta-binomial distributions reflect overdispersion, typical of aRME or X-inactivation, where allelic silencing varies stochastically across cells or tissues. Log-likelihood ratio tests were used to compare models, and statistical significance was assessed via one-tailed chi-squared tests with FDR correction (q ≤ 0.05), using the R packages VGAM (v1.1.12; Yee, 2010, 2020) and qvalue (v2.34.0; Storey et al., 2023) (**Sup Figure 4A**).

Within each subject, genes with adjusted p-values > 0.05 (i.e., following a binomial distribution) were further classified into the remaining allele expression types using R. This classification was based on three metrics: (i) the mean Allele 1 Ratio across tissues, (ii) the number of tissues with monoallelic expression, and (iii) the number of tissues with biallelic expression (**Sup Figure 4A**). Monoallelic expression was defined as an Allele 1 Ratio in the inclusive range of 0-0.05 or 0.95-1, and biallelic expression as a ratio in the exclusive range of 0.05-0.95. Constitutive monoallelic or biallelic expression refers to a consistent expression pattern across all tissues, whereas tissue-specific monoallelic expression is characterized by a mix of monoallelic and biallelic expression depending on the tissue. To further characterize biallelic expression, genes with a mean Allele 1 Ratio across tissues in the inclusive range of 0.45-0.55 were classified as showing balanced expression, while those with a mean ratio in the exclusive ranges of 0.05-0.45 or 0.55-0.95 were classified as showing biased expression.

To validate our approach, we selected genes with known allele expression patterns (**Sup Figure 4B-4C**). For biallelic balanced expression, we used *MAPK1* and the housekeeping genes *HNRNPA2B1*, *RAB11B*, *CSNK2B*, and *RHOA*; for biallelic biased, *ARL17A*; for tissue-specific imprinting, *LPAR6* and *UBE3A*; for constitutive imprinting, *NAP1L5* and *SNRPN*; and for aRME, *PARD6G*, *GALNT2*, *PIGG*, and *A4GALT* (Baran et al., 2015; Hounkpe et al., 2021; Kravitz et al., 2023; Pandey & Williams, 2015).

To assess the relationship between tissue involvement (affected vs. unaffected) and either gene expression or minor allele frequency (MAF) of CDG-causative genes, we conducted Spearman’s rank correlation analysis using GTEx data. Statistical significance was determined using a false discovery rate (FDR)-adjusted p-value threshold of ≤ 0.05. Correlation was considered biologically meaningful if the absolute value of Spearman’s coefficient (r) was ≥ 0.3.

### 2.5. Assessing Expression Quantitative Trait Loci (eQTLs) of CDG-causative genes

To assess the impact of eQTLs on CDG-causative gene expression levels, we quantified the number of eQTLs with positive and negative normalized effect sizes (NES) across 36 healthy GTEx tissues. To characterize eQTLs of CDG-causative genes, we extracted SNP ID data linked to eQTLs for each CDG-causative gene individually from Ensembl’s Human Short Variants (SNPs and indels, excluding flagged variants) and mapped to Ensembl Human Genes and Ensembl Human Regulatory Features using BioMart (GRCh38.p14, accessed August 2, 2024). To standardize annotation, eQTLs with multiple variant consequences (e.g., intron variant and splice polypyrimidine tract variant) were assigned the broader term (e.g., intron variant), and duplicates were removed. For eQTLs overlapping multiple regulatory regions, regulatory feature IDs and feature type descriptions were aggregated into single entries. Clinical significancy was evaluated through ClinVar (July 6, 2024) and the NHGRI-EBI Catalog of GWAS Associations v1.0.2 (gwas_catalog_v1.0.2-associations_e112_r2024-07-27.tsv).

### 2.6. Assessment of Immune Cell Composition and CDG-Causative Gene Expression Across Healthy Tissues

To assess the association between immune cell composition and either gene expression or minor allele frequency (MAF) of CDG-causative genes, we performed Spearman’s rank correlation analysis across 36 healthy tissues using data from the GTEx project. Immune cell composition estimates from the adult GTEx dataset were derived through RNA-based deconvolution, as described by Sobral et al. (2022). The cell types included B cells, M1 and M2 macrophages, monocytes, myeloid dendritic cells, neutrophils, natural killer (NK) cells, non-regulatory CD4⁺ T cells, CD8⁺ T cells, and regulatory CD4+ T cells (Tregs).

Statistical significance was defined as an adjusted p-value ≤ 0.05 using the false discovery rate (FDR) correction. Correlations were considered biologically meaningful when the absolute Spearman correlation coefficient (r) was ≥ 0.3.

### 2.7. Data Visualization

All plots were generated with R packages. Heatmaps were created with corrplot (v0.95) (Wie & Simko, 2024). Violin plots and stacked bar chart were produced using ggplot2(v3.5.1) (Wickham, 2016). The ggpubr (v0.6.0) (Kassambara, 2023), gridExtra (v2.3) (Auguie, 2017), and grid (v4.3.3) (R Core Team, 2024) packages were used for plot arrangement and annotations. Ridgelineplot was made using the ggplot2 (v3.5.1) (Wickham, 2016) and ggridges (v0.5.6) (Wilke, 2025). Sankey plots were generated with ggplot2 (v3.5.1) (Wickham, 2016) and ggsankey (v0.0.99999) (Sjoberg, 2024). Cell-specific dot plot showing gene expression across tissues were created using the DotPlot () function from the Seurat (v5.1.0) (Hao et al., 2024). Scatterplots were generated with base R’s plot() function.

### 2.8. Statistical Analyses and Code

Bioinformatic analyses were conducted in R (v4.3.3), Python (v3.12.3) and GNU Bash (v5.2.21). All code and supplementary data will be made available upon publication in a journal.

## Results

### 3.1 Tissue-Specific and Inter-Individual Variability in Expression of CDG-Causative Genes Reflects Phenotypic Heterogeneity

To investigate whether the phenotypic heterogeneity observed in CDG is reflected in the expression patterns of their causative genes, we analyzed transcriptomic data from 2,833 samples spanning 36 healthy adult human tissues from the GTEx project (Aguet et al., 2017; Carithers et al., 2015; Castel et al., 2020; Consortium et al., 2020; Eraslan et al., 2022) (**Sup Table 2**). All CDG-causative genes were expressed across different healthy tissues, although with varying expression levels. Notably, while most CDG types present neurological and muscular involvement, none of the causative genes showed peak expression in brain cortex or skeletal muscle (**Figure 1A, Sup Figure 1-2**). Gene expression across tissues also revealed inter-individual variability (**Sup Figure 2**), which may contribute to the phenotypic diversity observed in CDG.

**Figure 1.**
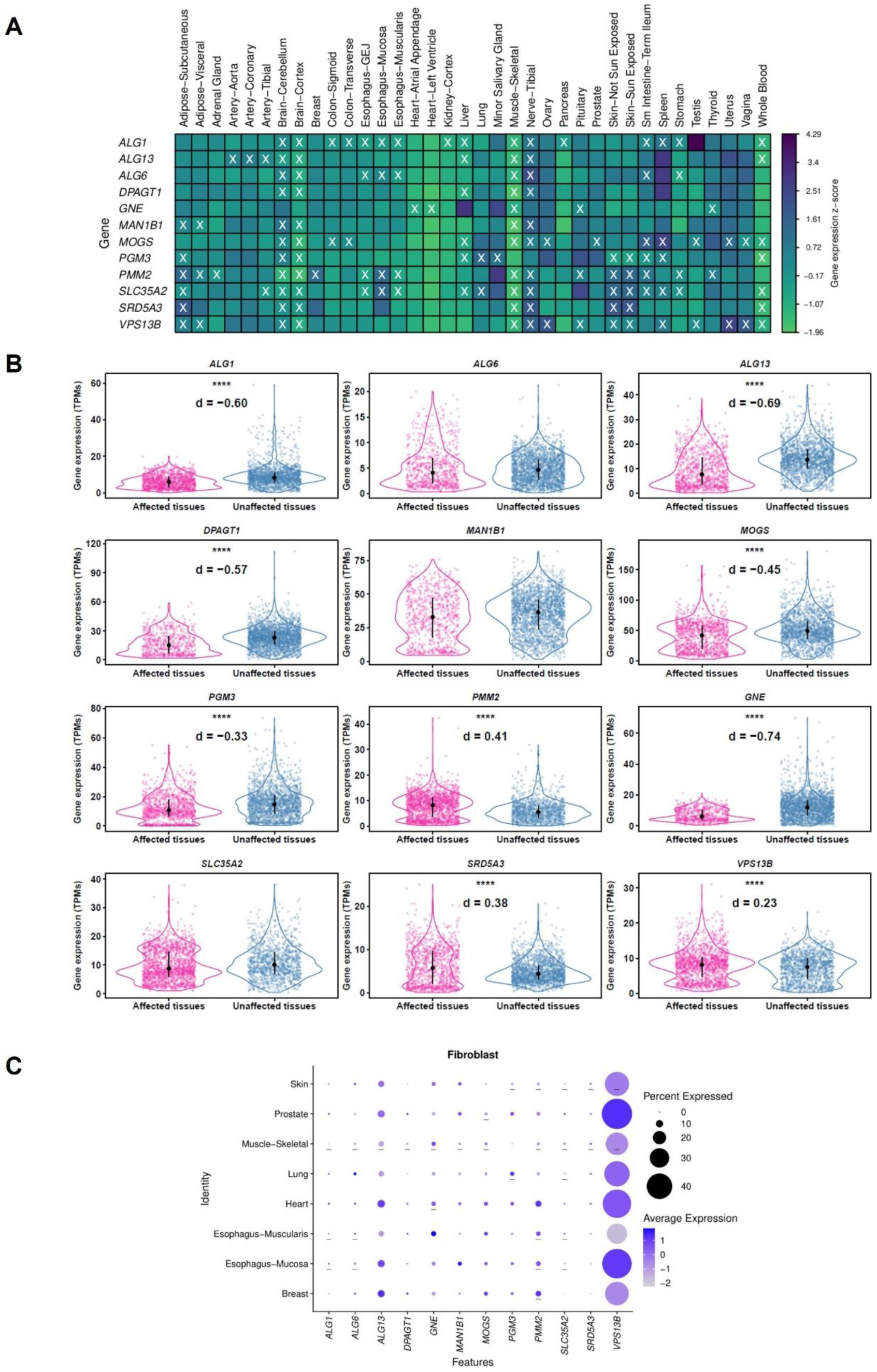
Expression of CDG-causative genes across healthy tissues. (A) Heatmaps with the expression profiles of CDG-causative genes across various healthy tissues (z-score of the median gene expression in Transcripts Per Million - TPM). The ‘x’ symbols highlight tissues most frequently affected by CDG types associated with these genes. (B) Violin plots comparing expression levels of CDG-causative genes between tissues clinically affected in CDG and those not affected. The affected tissues considered in this analysis are highlighed in figure 1A. The black dot in the middle represents the median and the thick black bar in the center represents the interquartile range. Asterisks indicate significant differences (****: *p* ≤ 0.0001). The letter *d* marks comparisons with effect size (Cohen’s *d*) > |0.2|. (C) Dotplot with the expression of 12 CDG-causative genes in fibroblasts across 8 healthy tissues. Dot size represents the percentage of cells expressing each gene, while color intensity reflects the log-normalized mean expression level among those cells expressing the gene. The ‘_’ symbols highlight tissues most frequently affected by CDG types associated with these genes.

**Figure 2.**
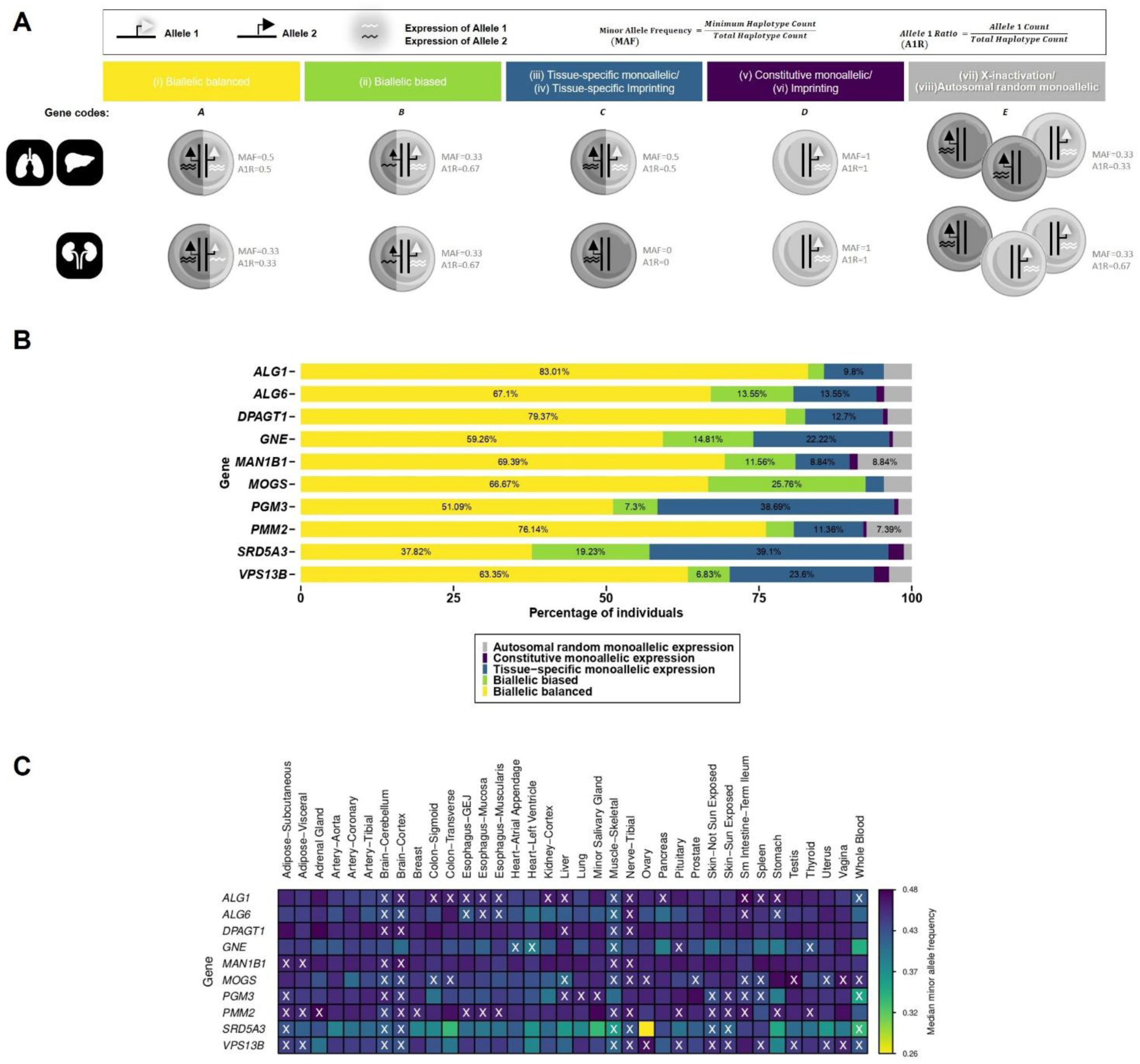
Allelic expression of CDG-causative genes across healthy tissues. (A) Schematic overview illustrating allele expression types in bulk RNA-seq data across a minimum of three tissues from an individual. The depicted cell or set of cells represents typical expression in the tissue. The overview provides information on expected allele counts, minor allele frequency (MAF) and allele 1 ratio (A1R) for genes with distinct allele expression profiles: Biallelic balanced expression; Biallelic biased expression; Tissue-specific monoallelic expression; Constitutive monoallelic expression; Tissue-specific imprinting or Imprinting. (B) Stacked barchart showing the percentage of individuals with different types of allele expression for each CDG-causative gene. Each type of allele expression is represented by a specific color. (C) Heatmap depicting the median minor allele frequency of CDG-causative genes across various healthy tissues. The ‘x’ symbols highlight tissues most frequently affected by CDG types associated with these genes.

Among the genes causing the most prevalent CDG types with widespread systemic involvement, including PMM2-, ALG1-, ALG13-, ALG6-, DPAGT1-, MAN1B1-, SLC35A2- and SRD5A3-CDG, several genes displayed tissue-specific expression peaks (**Figure 1A, Sup Figure 2**). For example, *ALG1* was highly expressed in the testis, while both *ALG13*, *ALG6* and *DPAGT1* showed elevated expression in the spleen. Interestingly, *SRD5A3* and *ALG6* exhibited moderately high expression in tissues commonly affected in their respective disorders, skin and tibial nerve, respectively. Notably, *PMM2*, the gene causing the most frequently diagnosed CDG, exhibited markedly high expression in the minor salivary glands, a tissue not typically associated with PMM2-CDG symptoms, suggesting a potentially overlooked role in disease pathophysiology. Interestingly, when contrasting affected and unaffected tissues across this CDG group, we found that *PMM2*, and *SRD5A3* were significantly upregulated in affected tissues, whereas *ALG1*, *ALG13*, and *DPAGT1* were significantly downregulated (**Figure 1B**).

*GNE* when defective causes a CDG (GNE myopathy) which is not systemic and primarily affects skeletal muscle. *GNE* expression was unexpectedly low in muscle and highest in the liver (**Figure 1A, Sup Figure 2**), indicating that tissue vulnerability may not always correlate with total gene expression levels. In agreement, total *GNE* expression is significantly elevated in tissues not affected by its defect (**Figure 1B**).

We also analysed the expression of the causative gene of CDGs characterized by major immune system involvement, MOGS-, PGM3- and VPS13B-CDG, to explore possible overlaps in pathophysiology and clinical presentation. Notably, *MOGS* and *PGM3* genes showed higher expression in tissues commonly affected in their respective disorders, namely the spleen and lung (**Figure 1A, Sup Figure 2**). However, these immune-related genes were not prominently expressed in the blood, and only *MOGS* showed high expression in spleen. Regarding other tissues where immune cells also significantly function, such as the lungs, gut, and skin (Farber, 2021), *PGM3* showed high expression in lung *and MOGS in the small intestine*. When contrasting affected and unaffected tissues within the immune-related CDG group, we found significant differences in total gene expression: *VPS13B* was elevated in affected tissues, whereas *MOGS* and *PGM3* were significantly reduced (**Figure 1B**).

Given that skin fibroblasts have been used as models for studying CDG (Gallego et al., 2024; Jabato et al., 2021; Lecca et al., 2011a; Lee et al., 2020; Pascoal et al., 2025; Shnaider et al., 2023), we also assessed single-cell expression profiles of the CDG-causative genes in fibroblasts from eight healthy tissues (GTEx). Among the genes analyzed, *VPS13B* showed the highest percentage of expressing fibroblasts across the eight tissues tested (**Figure 1C**). Its average expression level was also elevated in most fibroblast populations, particularly in the prostate and esophagus mucosa, with the notable exception of the esophagus muscularis. From the remaining CDG-causative genes, *ALG13*, *PMM2*, and *GNE* showed higher expression in fibroblasts from specific tissues. In skin, the top five CDG-causative genes with the highest percentage of expressing fibroblasts were *ALG13*, *GNE*, *MAN1B1*, *PMM2 and VPS13B.* Such findings support the use of fibroblasts as valuable models for exploring disease mechanisms, functional consequences of gene variants, and potential therapeutic targets in CDG-related pathophysiology.

Together, these findings reveal tissue-specific and inter-individual variability in expression of CDG-causative genes, offering mechanistic insights into the diverse clinical manifestations observed in CDG patients.

### 3.2. Allele-Specific Expression Patterns of CDG Causative Genes Revealed Tissue-Specific and Inter-Individual Variability

Given that allele-specific gene expression can influence disease manifestation, we conducted a comprehensive assessment of how the two alleles of the 12 CDG-causative genes are expressed across 36 healthy tissues using GTEx transcriptome data. Two complementary metrics were applied: allele 1 ratio (A1R) and minor allele frequency (MAF).

Based on A1R, we classified allele-specific expression of CDG-causative genes from each individual into five categories: (i) biallelic balanced, (ii) biallelic biased, (iii) tissue-specific monoallelic, (iv) constitutive monoallelic, and (v) autosomal random monoallelic expression (aRME) (**Figure 2A**). We then quantified the frequency of individuals assigned to each category. This approach was validated using control genes with well-established allele expression patterns (**Sup Figure 4B-4C**). The CDG-causative genes demonstrated diverse allelic expression profiles across all five categories (**Figure 2B, Sup Table 4**). A high proportion of individuals showed biallelic balanced expression, particularly for *ALG1* (83.01%), *DPAGT1* (79.37%), and *PMM2* (76.14%). In contrast, biallelic biased expression was more frequently observed for *MOGS* (25.76%) and *SRD5A3* (19.23%). Several genes also showed a high proportion of tissue-specific monoallelic expression, especially *SRD5A3* (39.10%), *PGM3* (38.69%), *VPS13B* (23.60%), and *GNE* (22.22%), which may help explain the variability in clinical symptoms. Autosomal random monoallelic expression was observed in a smaller subset of individuals, most notably in *MAN1B1* (8.84%) and *PMM2* (7.39%). Constitutive monoallelic expression was rare across all genes.

Across 36 healthy tissues, the median minor allele frequency (MAF) of CDG-causative genes ranged from 0.26 to 0.48 (**Figure 2C, Sup Table 2**), indicating that allelic expression was not always strictly equal between the two alleles. Nevertheless, balanced biallelic expression (MAF ∼0.5) remained the predominant pattern. Several genes, most notably *SRD5A3*, *GNE*, *PGM3*, and *VPS13B*, displayed marked reductions in MAF, particularly in tissues clinically affected by their respective CDG types. For *SRD5A3*, the lowest MAF was observed in the ovary (0.26), with additional reductions in the transverse colon and minor salivary gland (both 0.33), the stomach (0.35), whole blood (0.33), and skeletal muscle (0.36). *GNE* showed decreased MAF in the left ventricle (0.39) and blood (0.35), while *PGM3* exhibited lower values in blood (0.35) and stomach (0.39). Similarly, *VPS13B* presented reduced MAF in the stomach and left ventricle (both 0.38).

These patterns suggest that allelic imbalance is not stochastic but enriched in tissues most vulnerable to disease. These findings also highlight substantial inter-individual and tissue-specific variability in allelic expression of CDG-causative genes. Such regulatory diversity may contribute to the variable penetrance and expressivity observed in CDG patients.

### 3.3. Tissue-Specific eQTLs Reveal Regulatory Basis of Phenotypic Variability in CDGs

While CDGs are primarily diagnosed based on coding mutations in causative genes, regulatory variants, such as intronic and intergenic SNPs, can also modulate gene expression as expression quantitative trait loci (eQTLs) and contribute to disease heterogeneity. To investigate this regulatory dimension, we used the GTEx atlas of genetic regulatory effects (Consortium et al., 2020) to assess eQTLs for CDG-causative genes and identified the tissues in which their effects are most pronounced. We quantified these effects using the normalized effect size (NES) of each eQTL, a metric that indicates whether a variant increases (positive NES) or decreases (negative NES) gene expression, and by how much.

The number of unique eQTLs affecting gene expression varied considerably among CDG-causative genes, ranging from only 21 eQTLs identified for *SLC35A2* to 1232 for *SRD5A3* (**Figure 3A**). Interestingly, while certain genes such as *MOGS*, and *SRD5A3* displayed a higher proportion of eQTLs associated with increased expression, others like *MAN1B1* and *GNE* were predominantly associated with decreased expression **(Figure 3A)**. Tissue-specific analysis revealed distinct patterns of regulatory impact. Some CDG-causative genes, such as *MAN1B1*, *MOGS*, *SRD5A3* and *PMM2*, exhibited widespread regulatory effects across multiple tissues, whereas others displayed eQTLs with more tissue-restricted activity (e.g., *ALG13*, *DPAGT1*, *PGM3*, *SLC35A2* and *VPS13B*) (**Figure 3B**).

**Figure 3.**
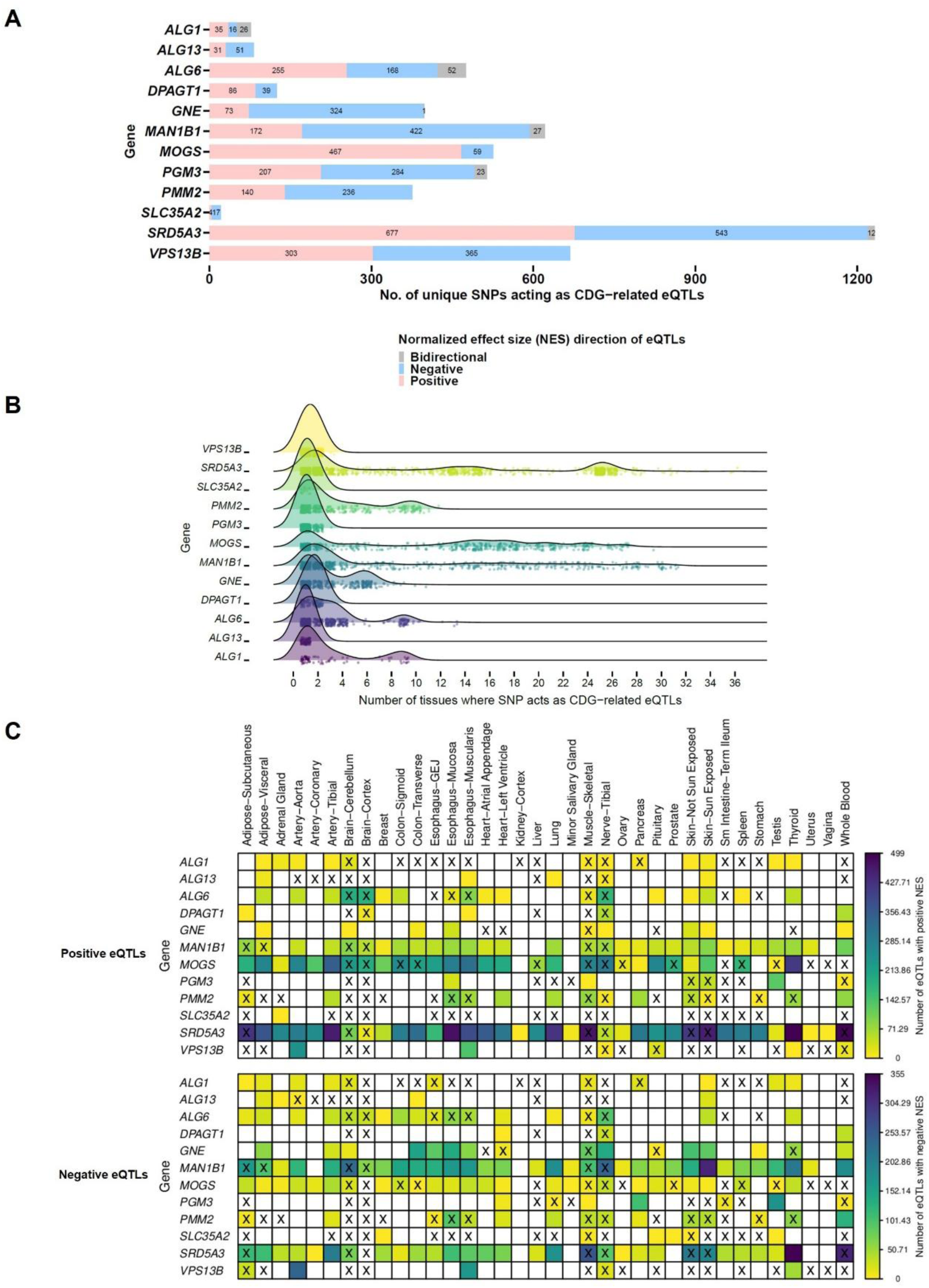
eQTLs for CDG-causative genes across healthy tissues. (A) Stacked barplot showing the number of unique SNPs acting as eQTLs with positive, negative, or bidirectional NES across tissues. (B) Ridgeline plot depicting the number of tissues harboring the same eQTL for CDG-causative genes. (C) Heatmaps showing the number of eQTLs with positive and negative NES for CDG-causative genes across healthy tissues. The ‘x’ symbols highlight tissues most frequently affected by CDG types associated with these genes.

Furthermore, several genes showed both positive and negative eQTLs in tissues frequently implicated in their clinical phenotypes (**Figure 3C**). For instance, both upregulating and downregulating eQTLs were found for *PMM2* in adipose-subcutaneous tissue, esophagus, skeletal muscle, tibial nerve, skin, stomach, and thyroid, and for *SRD5A3* in subcutaneous adipose tissue, cerebellum, skeletal muscle, tibial nerve, and skin.

To examine the clinical relevance of these regulatory variants, we queried the NHGRI-EBI GWAS Catalog (Cerezo et al., 2025). A significant proportion of the identified eQTLs were linked to disease-associated traits (**Sup Table 6**). Notably, negative eQTLs in *PMM2* were associated with reduced levels of the encoded protein, phosphomannomutase 2 (SNP ID: rs11645498, rs3826198 and rs78918345), and with gastroesophageal reflux disease (SNP ID: rs2270288). For *SRD5A3*, several eQTLs were linked to traits such as subcutaneous adipose tissue composition (SNP ID: rs10001607, rs10462028, rs11133373, rs11935857, rs13132420, rs13142096, rs13434995, rs1873091, rs34929896, rs3805383, rs3805389, rs4455482, rs55796004, rs57826934, rs62303684, rs62303689, rs6554274, rs6840236, rs7663650, rs861029), blood cell parameters (SNP ID: rs13127906, rs13139100, rs34658078), brain morphology (SNP ID: rs34658078), and skin characteristics (SNP ID: rs181411540 and rs11724094).

Together, these findings suggest that regulatory variation of tissue-specific and directionally diverse eQTLs may contribute significantly to the phenotypic heterogeneity observed in CDG.

### 3.4. CDG-Related eQTLs Are Predominantly Enriched in Intronic Enhancers of Distinct Genes

To further characterize the regulatory and functional context of eQTLs affecting CDG-causative genes, we examined their NES direction across tissues (positive, negative and bidirectional) and genomic location, assessing whether they occur within CDG-causative genes, other genes, or in intergenic regions. We also evaluate their overlap with regulatory elements (open chromatin regions, transcription factor binding sites, predicted promoters, enhancers, and CTCF binding sites) (**Sup Table 6**). Overall, eQTLs with negative NES were slightly more common than those with positive NES, while bidirectional eQTLs were relatively rare (**Figure 4A, Sup Figure 5**). Among the 12 CDG-causative genes, we identified 1235 eQTLs overlapping with at least one regulatory element. *MOGS* and *SRD5A3* exhibited the highest number of positive eQTLs in open chromatin region, promoters and enhancers, while *MAN1B1* and *SRD5A3* led for negative eQTLs (**Figure 4A**). Bidirectional eQTLs intersecting multiple regulatory regions were primarily observed in *MAN1B1* (**Sup Figure 5**).

**Figure 4.**
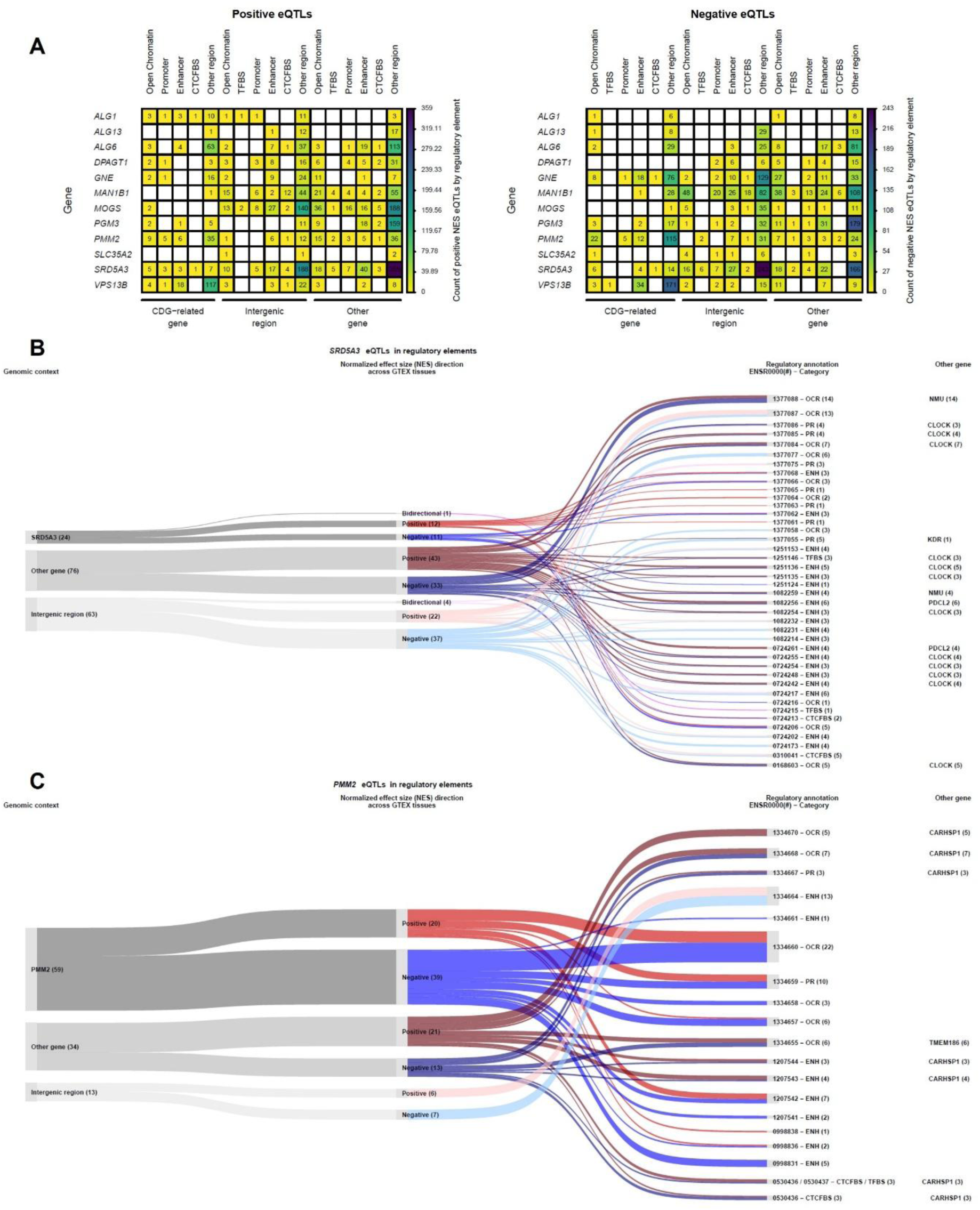
Genomic Location of CDG-related eQTLs. Regulatory categories include open chromatin regions (OCR), transcription factor binding sites (TFBS), predicted promoters (PP), predicted enhancers (ENH), CTCF binding sites (CTCFBS), and other regions. (A) Heatmaps showing the number of eQTLs with positive or negative NES located in regulatory regions of the gene itself, in intergenic regions, or within another gene. (B-C) Sankey plots illustrating the number and distribution of eQTLs for the *PMM2* (B) and *SRD5A3* (C) genes across regulatory regions. Displayed regulatory regions were selected based on their location and eQTL count: those overlapping the gene itself include ≥1 eQTL, while those overlapping another gene and/or an intergenic region include ≥3. Flows connect three eQTL classifications (nodes): (1) Genomic context - intergenic, within the gene itself, or another gene (grey shades); (2) NES direction across GTEx tissues - positive (red shades), negative (blue shades), or bidirectional (purple shades); (3) Regulatory annotation (Ensembl ID and category). Flow shading reflects genomic context: light for intergenic regions, medium for eQTLs within the gene itself, and dark for those within another gene. The final column lists the names of other genes, where applicable.

**Figure 5.**
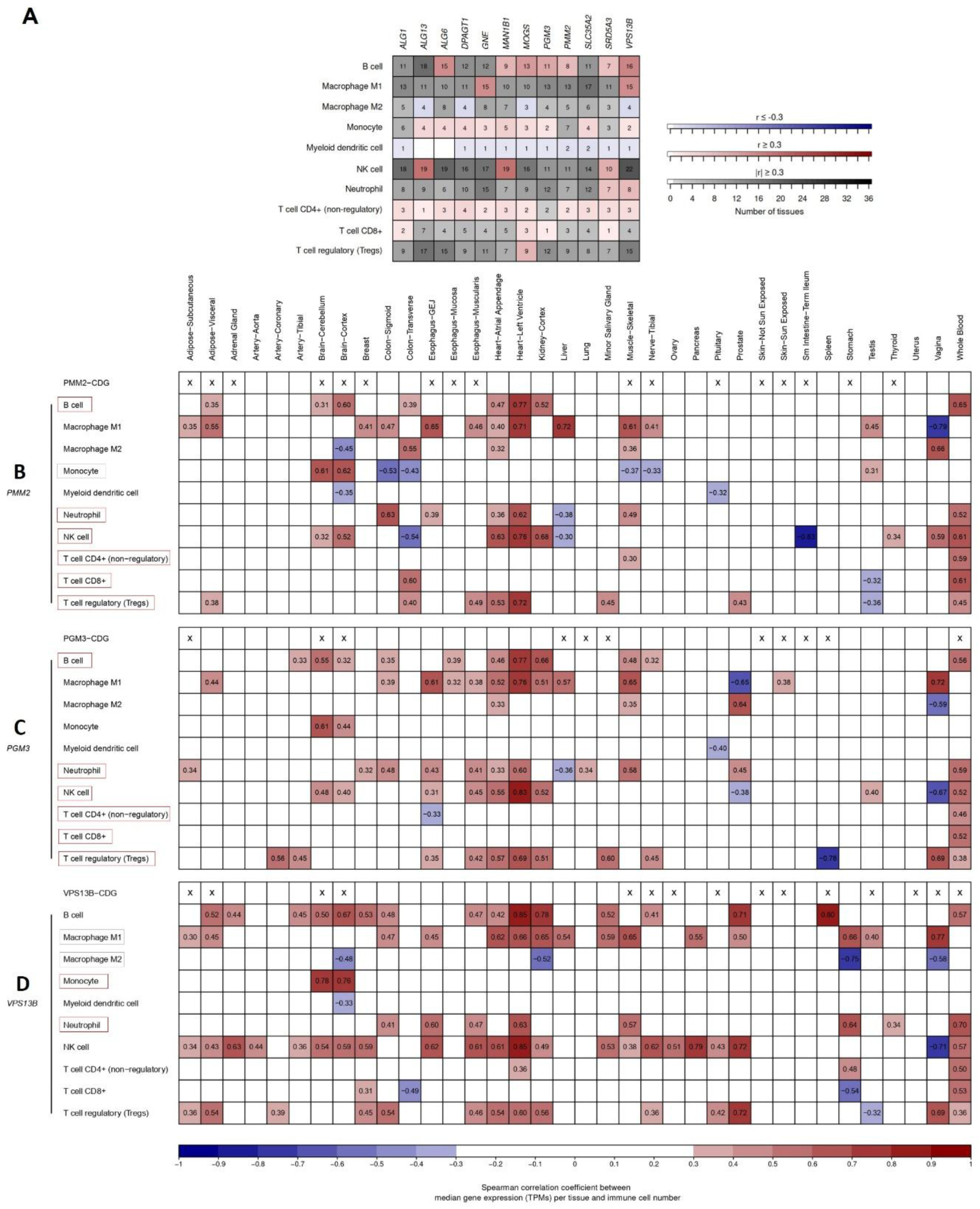
Immune cell types associated with CDG-causative gene expression. (A) Heatmap summarizing the number of tissues with statistically and biologically significant correlations between gene expression and immune cell abundance using RNA-based deconvolution (FDR-adjusted *p* ≤ 0.05 and |Spearman r| ≥ 0.3). Color indicates correlation sign: all positive (red), all negative (blue) and positive and negative (grey). White indicates no significant correlation. (B-D) Heatmaps showing significant correlations between median gene expression (TPMs) of *PMM2*, *PGM3*, and *VPS13B* and immune cell counts across tissues (FDR-adjusted *p* ≤ 0.05 and |Spearman r| ≥ 0.3). Correlation strength is indicated by color intensity for positive (red) and negative correlation (blue). White indicates no significant correlation. An ‘x’ in the top row marks tissues frequently affected in CDG types associated with each gene.

Strikingly, the majority of eQTLs were not located within the CDG-causative genes they regulate but instead resided in other genes or intergenic regions (**Sup Table 6**). Among all regulatory elements, predicted enhancers were the most frequently associated with CDG-related eQTLs, regardless of the direction of effect (**Figure 4A**). Enhancers embedded within other genes accounted for the highest number of eQTLs for *SRD5A3*, *PGM3*, *MOGS*, *ALG6* and *MAN1B1* (Figure 4A-4C, Sup Figure 6). Notably, *SRD5A3* eQTLs were frequently mapped to *CLOCK* (a transcription factor involved in circadian rhythm), *PDCL2* (a putative modulator of heterotrimeric G proteins essential for male fertility), *NMU* (a neuropeptide linked to circadian rhythm and other processes), and *SRD5A3-AS* (a long non-coding RNA antisense to *SRD5A3*), with strong enrichment in enhancer regions (Figure 4B). Likewise, gene-level mapping revealed that most *PMM2* eQTLs were located within the *CARHSP1* gene encoding an RNA binding protein and overlapped with all five regulatory element types, particularly predicted enhancers (Figure 4C). Consistent with the predominance of enhancer-associated eQTLs, a large proportion of variants were in intronic regions (**Sup Figure 7, Sup Table 6**). For positive NES eQTLs, *SRD5A3*, *MOGS*, and *PGM3* were most frequently found in the introns of other genes, whereas *VPS13B* had the highest number within its own introns. Negative NES eQTLs were most common in the introns of other genes for *PGM3*, *SRD5A3*, and *MAN1B1*, and within self-introns for *VPS13B* and *PMM2*. Bidirectional eQTLs were primarily located in introns of other genes for *ALG6* and *PGM3*.

Together, these findings highlight that CDG-related eQTLs are predominantly localized in intronic enhancer regions of genes other than the ones they regulate. This underscores a complex, tissue-specific regulatory landscape that likely contributes to the variable expression patterns and clinical heterogeneity observed in CDG.

### 3.5. Immune Cell Profiles Are Associated with CDG-Causative Gene Expression Across Various Tissues

To assess the influence of CDG-causative genes on the immune microenvironment, we analyzed RNA-based immune cell deconvolution data from healthy tissues in the GTEx dataset (Sobral et al., 2022). Specifically, we examined correlations between CDG gene expression and the abundance of immune cell types across multiple tissues (**Sup Table 7**).

All 12 CDG-causative genes exhibited statistically significant correlations with various immune cell types, predominantly positive (**Figure 5A, Sup Table 7**). Interestingly, B cells displayed several unique positive correlations with the expression of seven CDG-causative genes across multiple tissues *(ALG6, MAN1B1, MOGS, PGM3, PMM2, SRD5A3* and *VPS13B).* M1 macrophages, NK cells, and Tregs also showed distinct positive correlations with specific CDG-causative genes across several tissues. Non-regulatory CD4⁺ T cells and monocytes tended to correlate positively with most CDG-causative genes, while myeloid dendritic cells generally showed negative correlations, though both patterns were limited to a smaller number of tissues. For specific CDG-causative genes, M1 macrophages exhibited exclusively unique positive correlations, whereas M2 macrophages were associated with unique negative correlations across tissues. Notably, both NK cells and M1 macrophages demonstrated a mix of positive and negative correlation patterns with CDG-causative gene expression across multiple tissues, underscoring their potential relevance to tissue-specific gene regulation in CDG.

Notably, most genes showed stronger correlations with B cells, NK cells, and M1 macrophages in the left ventricle of the heart (**Figure 5B-5D, Sup Figure 11-12, Sup Figure 8**). Although the heart is not typically affected in the CDG subtypes studied, this finding may indicate tissue-specific regulatory interactions or underexplored systemic effects. Additionally, opposing correlations were observed for M1 and M2 macrophages with specific CDG-causative genes in tissues such as the kidney cortex, stomach, and vagina (**Figure 5B-5D, Sup Figure 11-12, Sup Figure 9**), suggesting differential immune modulation across environments.

We then focused on five genes with known or suspected immune involvement in CDG, *PMM2, PGM3, VPS13B, MOGS* and *SLC35A2,* analyzing their expression correlations with immune cells across tissues commonly affected in their respective CDG types. Blood was included for its immunological relevance, even in cases like PMM2-CDG, where it is not typically affected.

*PMM2* expression positively correlated with B cells in visceral adipose tissue, brain regions (r = 0.31-0.60), and blood (r = 0.65), aligning with reports of B cell expansion in PMM2-CDG (Blank et al., 2006). Neutrophils showed positive associations in the gastroesophageal junction, skeletal muscle, and blood (r = 0.52), consistent with reported neutropenia (Monin et al., 2014). NK cells were positively associated in brain, thyroid, and blood (r = 0.61), and negatively in the terminal ileum (r = −0.83) (**Figure 5B, Sup Figure 10**), echoing observations of increased NK cell counts in severe cases (García-López et al., 2016). All T cell subsets correlated positively with *PMM2* in blood (r = 0.45-0.61), with additional associations for non-regulatory CD4⁺ T cells in skeletal muscle and Tregs in adipose tissue and esophagus. Notably, severely affected patients showed increased NK cells, marked CD8⁺ T cell reductions, and transient CD4⁺ T cell decline (García-López et al., 2016). Monocytes correlated positively in the brain (r = 0.61-0.62), negatively in skeletal muscle and tibial nerve, and showed no correlation in blood (**Figure 5B)**, consistent with findings from de Haas et al. (2022).

*PGM3* expression correlated with B cells in blood (r = 0.56) and brain (r = 0.32-0.55), and with neutrophils in blood, lung, and adipose tissue (r = 0.59) (**Figure 5C, Sup Figure 10**), consistent with reduced CD27⁺ memory B cells and neutropenia (Yang et al., 2024). Positive correlations with non-regulatory CD4⁺ T cells and Tregs in blood and salivary gland (r = 0.60), and a negative one in spleen (r = −0.78), mirror clinical findings of CD4⁺ T cell reduction and impaired Treg differentiation (Yang et al., 2024). Expression also correlated with CD8⁺ T cells and NK cells in blood (r = 0.52), and NK cells in brain (r = 0.40-0.48), reflecting their variable levels in patients (Pascoal et al., 2024).

*VPS13B* expression showed strong correlations with neutrophils in blood (r = 0.70) and skeletal muscle (r = 0.57), and with monocytes in brain (r = 0.76-0.78) (**Figure 5D, Sup Figure 10)**, in line with observed neutropenia and altered monocyte counts in VPS13B-CDG (Pascoal et al., 2024). Interestingly, the minor allele frequency of *VPS13B* was strongly negatively correlated with M1 macrophages in the terminal ileum (r = −1), representing the only significant immune correlation linked to genetic variation (**Sup Figure 13, Sup Table 8**).

*MOGS* gene expression was positively associated with B cells in the brain, intestine, testis, and blood (r = 0.72), and with T cells in blood (r = 0.47-0.72) and terminal ileum Tregs (r = 0.83) (**Sup Figure 11**). These results reflect the B and T lymphocytosis, T lymphopenia, and reduced lymphocyte proliferation observed in MOGS-CDG (Pascoal et al., 2024).

Neutrophil correlations in skeletal muscle and blood (r = 0.59) support reported variability in neutropenia and neutrophilia (Pascoal et al., 2024).

*SLC35A2* expression correlated with non-regulatory CD4⁺ T cells in blood (r = 0.39) and with NK cells in lung and skin (**Sup Figure 12)**, reflecting its known role in glycosylation and immune regulation (Itell et al., 2024; Xu et al., 2023).

These results collectively underscore the significant impact of CDG-causative genes on immune cell regulation across multiple tissues, revealing important insights into the immunopathology of CDGs.

## Discussion

Given the limited understanding of phenotypic variability in CDGs and the difficulty of obtaining biopsies from affected tissues, we systematically assessed the expression and regulatory landscape of CDG-causative genes using transcriptome data from healthy adult GTEx tissues.

Gastrointestinal symptoms like vomiting in infancy are often under-reported in CDG, partly because they are non-specific and can arise from various causes (Kamarus Jaman et al., 2021). Similarly, in our tissue-specific symptom assignments, we treated vomiting as non-specific, which may have contributed to underestimating gastrointestinal involvement.

Our analysis revealed that CDG-causative genes are broadly but heterogeneously expressed across tissues, offering insights into the tissue-specific symptoms observed in various CDG types. In many cases, elevated gene expression aligned with clinical features, for instance, *SRD5A3* in skin, *MOGS* in spleen, and *ALG6* in peripheral nerves. Interestingly, *PMM2* showed strong expression in minor salivary glands, suggesting a potential molecular basis for the feeding difficulties observed in PMM2-CDG (Oliveira et al., 2025). Likewise, *VPS13B* exhibited sex-specific expression in reproductive tissues, which may relate to the delayed puberty characteristic of its associated disorder (Momtazmanesh et al., 2020). Importantly, low expression in a tissue did not preclude involvement in disease, as seen in blood, brain, and muscle. These discrepancies could reflect heightened glycosylation demands, limited redundancy, or developmental expression windows. Supporting this idea, only combined prenatal and postnatal sialic acid treatment improved neurodevelopmental outcomes in NANS-CDG (den Hollander et al., 2021). Additionally, a tissue-specific RNA variant generated by alternative splicing may underlie the pathology of a given tissue, rather than its total gene expression. For example, Awasthi et al. (2022) linked skeletal muscle pathology in GNE-CDG(ar) to a muscle-specific isoform of *GNE*. In addition, gene expression across tissues revealed inter-individual variability, which may contribute to the phenotypic diversity observed in CDG.

To complement tissue-wide expression patterns, we next examined gene expression in fibroblasts, which are widely used for disease modeling. Among the CDG genes, *VPS13B* emerged as the most broadly and consistently expressed across fibroblasts from eight different tissues. This supports its utility in fibroblast-based assays and aligns with previous studies using patient-derived fibroblasts to generate iPSC-derived neurons for detailed molecular analyses (Lee et al., 2020; Shnaider et al., 2023). In contrast, *ALG13*, *PMM2*, and *GNE* displayed more tissue-restricted expression, underlining the need to consider cellular context when interpreting disease models. Within skin-derived fibroblasts, six CDG genes (*VPS13B*, *ALG13*, *GNE*, *MAN1B1*, *PMM2*, *ALG6*) were highly expressed, and transcriptomic studies have already uncovered dysregulated pathways in several corresponding CDG types (e.g., DPM1-, ALG12-, ALG6-, and PMM2-CDG) (Gallego et al., 2024; Jabato et al., 2021; Lecca et al., 2011b; Pascoal et al., 2025), reinforcing the relevance of fibroblast models for specific CDG subtypes.

While total gene expression provides a foundational understanding, allelic expression analysis offers additional resolution into individual-level regulatory differences that may contribute to variable disease severity. Most CDG-causative genes showed balanced biallelic expression, as expected; however, specific deviations were observed. For example, *SRD5A3* displayed reduced MAF in ovaries, where symptoms such as ovarian insufficiency have been reported (Kamarus Jaman et al., 2021). Other tissues with reduced *SRD5A3* MAF, such as the salivary gland, stomach, and colon, also align with reported gastrointestinal symptoms in SRD5A3-CDG, including sucking and swallowing discoordination, GERD, and irritable bowel syndrome (Al-Gazali et al., 2008; Kamarus Jaman et al., 2021; Kasapkara et al., 2012; Morava et al., 2009, 2010; Tuysuz et al., 2016). Reduced *SRD5A3* MAF in blood and skeletal muscle may help explain the frequent occurrence of microcytic anemia (Garapati et al., 2024; Kara et al., 2014) and hypotonia (Holla et al., 2023; Jaeken et al., 2020; Kara et al., 2014) in SRD5A3-CDG. Similarly, *GNE*, *PGM3*, and *VPS13B* showed tissue-specific reductions in MAF that correspond with their known clinical features. For instance, *GNE* MAF was reduced in blood, consistent with its known role in thrombocytopenia, either isolated or in combination with GNE myopathy (Izumi et al., 2014; Jang et al., 2022; Revel-Vilk et al., 2018). Although platelets are anucleate, they can modulate gene expression in leukocytes (Franks et al., 2010). Low *GNE* MAF in the left ventricle may contribute to the reported cardiac involvement in GNE-CDG, including first-degree AV block, ST-T changes, signs of myocardial infarction, and sleep apnea syndrome (Mori-Yoshimura et al., 2022; Yoshioka et al., 2022). Reduced *PGM3* MAF in blood and stomach aligns with immune defects and gastrointestinal symptoms such as GERD, obstruction, malrotation, and recurrent infections (Jaeken et al., 2019; Pascoal et al., 2024; Yang et al., 2024). *VPS13B* showed low MAF in the stomach and left ventricle, consistent with feeding difficulties and cardiac anomalies (e.g., ventricular dysfunction) reported in VPS13B-CDG (Momtazmanesh et al., 2020).

To better characterize expression heterogeneity, we classified CDG-causative genes into five allelic expression types across tissues. While balanced biallelic expression was most common (particularly in *ALG1*, *DPAGT1*, and *PMM2*) biallelic biased expression was frequent in *MOGS* and SRD5A3, and tissue-specific monoallelic expression was notable in *SRD5A3*, *PGM3*, *VPS13B*, and *GNE*. Constitutive monoallelic expression was rare. These allele-specific patterns have been linked to cis-eQTLs (Kravitz et al., 2023) and are known to influence variable penetrance and expressivity in diseases such as pulmonary fibrosis and immune-related HLA conditions (Johansson et al., 2021, 2022; Natri et al., 2024; Yamamoto et al., 2020). Similarly, these allele expression types may help explain clinical variability in CDG. Atypical random allelic expression was rare but observed in *MAN1B1* and *PMM2*, and has been implicated in cancer, cardiovascular, and neurodevelopmental disorders, as well as inborn errors of immunity (Kravitz et al., 2023; Stewart et al., 2025). Although CDG-causative genes such as *PIGG*, *A4GALT*, and *GALNT2* showed high random allelic expression frequency, the ten genes profiled in this study displayed such type only in subsets of individuals, suggesting a potential but limited contribution to CDG phenotypic diversity.

To explore the genetic basis of regulatory variation, we examined eQTLs across CDG genes. The number and tissue distribution of unique eQTL SNPs varied widely, reflecting diverse regulatory landscapes. *SRD5A3*, *MOGS*, *MAN1B1*, and *VPS13B* had many eQTLs, but only the first three showed cross-tissue regulation. This contrast aligns with known differences in broad versus tissue-specific eQTL effects (Sul et al., 2013). As expected, X-linked genes (*SLC35A2*, *ALG13*) had fewer eQTLs due to underrepresentation of X chromosome regulatory variation (Kukurba et al., 2016). Directionality also varied: *MOGS* was predominantly influenced by positive NES eQTLs, *MAN1B1* by negative ones, and *SRD5A3* by both, often within clinically relevant tissues. These findings support the notion that regulatory variation shapes tissue-specific phenotypes in CDG.

Linking GTEx eQTLs to the NHGRI-EBI GWAS Catalog further validated their functional relevance. CDG-related eQTLs were associated with traits affecting tissues impacted by their corresponding diseases. For example, *PMM2* with brain and esophagus traits, and *SRD5A3* with adipose tissue, blood, brain, and skin. Notably, three *PMM2* eQTLs with negative eQTLs were linked to reduced protein levels, highlighting their translational potential. We then characterized the regulatory and genomic context of these eQTLs. While general GTEx eQTLs are often found in promoters and open chromatin (Consortium et al., 2020), CDG-related eQTLs were more commonly located in enhancers, particularly *MAN1B1*, *SRD5A3*, and *MOGS*. Many mapped outside the target gene, often to enhancers located in other genes or intergenic regions. For instance, several *PMM2* eQTLs were found in regulatory elements within *CARHSP1*, a gene involved in mRNA stabilization and transcriptional regulation (Hou et al., 2011; Li et al., 2016; Lindquist et al., 2014; Pfeiffer et al., 2011; Schäfer et al., 2003). Similarly, *SRD5A3* eQTLs mapped to enhancers in *CLOCK* and *NMU*, genes involved in circadian regulation (Hogenesch et al., 1997, 1998; Malendowicz & Rucinski, 2021; Martinez & O’Driscoll, 2015). Notably, *Srd5a3* displays 24-hour rhythmic expression in chicken pineal glands under both light-dark and constant darkness (Chustecka et al., 2021), suggesting it may be clock-controlled. These enhancer-associated eQTLs may mediate gene expression through long-range chromatin looping (Andersson & Sandelin, 2020; Krivega & Dean, 2012), and disruptions may underlie disease pathogenesis (Mitkin et al., 2018; Ustiugova et al., 2019; A. Uvarova et al., 2023; A. N. Uvarova et al., 2022). Supporting these findings, variant consequence analysis showed that CDG-related eQTLs were predominantly located in introns, frequently within other genes, reinforcing the importance of distal regulation. Exceptions included *VPS13B*, which had many eQTLs within its own introns, and *PMM2*, which showed this pattern for negative NES eQTLs.

To investigate immune involvement in CDG, we correlated CDG-causative gene expression with immune cell abundance across healthy GTEx tissues. Tissue-specific correlations, including opposing M1/M2 macrophage patterns, suggest diverse immune modulation. Although heart issues are rare in most CDG types (∼6%) (Zemet et al., 2024), several CDG genes showed notable immune correlations in the healthy heart. The hexosamine biosynthesis pathway (HBP), involving PGM3 and producing UDP-GlcNAc for N-glycosylation, N-glycan branching, and O-GlcNAcylation, plays a dual role in the heart: acute activation is cardioprotective during stress (e.g., ischemia/reperfusion), while chronic activation promotes hypertrophy (Gélinas et al., 2018; Tran et al., 2020; Wang et al., 2014). *PGM3* expression is elevated in hypertrophic heart tissue (Tran et al., 2020), though heat immune issues (e.g., pericarditis) are rare in PGM3-CDG (Fusaro et al., 2021). PMM2-CDG can involve isolated cardiac infections (e.g., pericarditis) and systemic infections with cardiac effects (e.g., pericardial effusion) (Blank et al., 2006; de Lonlay et al., 2001; Noelle et al., 2005; Truin et al., 2008).

While most immune data in CDG come from blood, our findings reinforce the role of blood-associated immune cells and suggest their activity in other affected tissues. In healthy blood, *PMM2* expression correlated with neutrophils, B, NK, and T cells, but not monocytes, echoing some PMM2-CDG reports of neutropenia, B lymphocytosis, elevated NK cells, reduced CD4⁺/CD8⁺ T cells, normal monocyte levels with glycosylation defects (Asteggiano et al., 2018; Blank et al., 2006; de Haas et al., 2022; García-López et al., 2016; Monin et al., 2014; Wu et al., 2018). These cells also correlated in GI tissues, brain, visceral adipose tissue, skeletal muscle, liver, and thyroid. PMM2-CDG has been linked to GI and brain inflammation, and infections often leading to neurological effects like seizures and stroke-like episodes (Arnoux et al., 2008; Brum et al., 2008; Francisco et al., 2020; García-López et al., 2016; Görlacher et al., 2020; Kiparissi et al., 2023; Serrano, 2021; Shanti et al., 2009; Tiwary et al., 2022).

*PGM3* expression correlated with neutrophils, B, T, and NK cells in healthy blood, consistent with PGM3-CDG features like neutropenia, reduced CD27⁺memory B and CD4⁺ T cells, and variable CD8⁺ T/NK cell levels (Pascoal et al., 2024; Yang et al., 2024). In vitro, PGM3 inhibition in stimulated CD4⁺ T cells promotes Th1/Th2 over Th17/Treg responses (Yang et al., 2024). These immune cells also correlated with *PGM3* in the brain, lung, subcutaneous adipose tissue, liver, minor salivary gland, and spleen, tissues commonly affected by immune symptoms in PGM3-CDG (Pascoal et al., 2024). Respiratory allergies, infections and bronchitis suggest lung dysfunction, while elevated IgE and eczema/dermatitis implicate both spleen and lung. Variable IgG levels may involve the liver, spleen and salivary gland. Skin infections and inflammation (e.g. folliculitis and dorsal skin inflammation) point to subcutaneous adipose tissue.

*VPS13B* expression correlated with neutrophils in blood and muscle, and with monocytes in the brain, reflecting VPS13B-CDG features such as frequent neutropenia, rare monocytopenia/monocytosis, and isolated brain infections like meningitis (Pascoal et al., 2024).

*MOGS* expression correlated with neutrophils, B, and T cells in blood, reflecting MOGS-CDG features including variable neutrophil counts, B/T lymphocytosis, T lymphopenia, and impaired lymphocyte proliferation (Pascoal et al., 2024). These patterns extended to brain, intestine, testis, and muscle. Immune involvement of the brain and intestine in MOGS-CDG is suggested by infections (e.g., meningitis, bacterial enteritis), food allergies, and low IgA/IgM levels (Pascoal et al., 2024).

Although SLC35A2-CDG is not strongly immune-related, *SLC35A2* expression correlated with non-regulatory CD4⁺ T cells in blood and NK cells in lung and skin. SLC35A2 inactivation alters HIV tropism via glycan truncation in CD4⁺ T cells (Itell et al., 2024), and *SLC35A2* expression inversely correlates with NK cell infiltration in lung and skin cancers (Xu et al., 2023), suggesting context-dependent immune roles.

*VPS13B* minor allele frequency was negatively correlated with M1 macrophages in the terminal ileum, consistent with gastrointestinal involvement in VPS13B-CDG, including neonatal feeding difficulties (Momtazmanesh et al., 2020) and a single report of inflammatory bowel disease (Pascoal et al., 2024).

Our integrative analysis of CDG-causative genes across healthy human tissues highlights the complex interplay between gene expression, allelic regulation, and genetic variation in shaping the phenotypic diversity of CDG. We demonstrate that while tissue-specific expression patterns partly explain clinical involvement, factors such allelic expression imbalance and tissue-specific eQTLs may contribute significantly to disease variability. The distinct regulatory landscapes we identified, including enhancer-associated eQTLs within introns of other genes and in intergenic regions, underscore the importance of considering distal regulatory elements in CDG pathogenesis. Furthermore, the consistent expression of key CDG genes in fibroblasts reinforces their utility as disease models, especially for mechanistic studies and therapeutic exploration. Correlations between gene expression and immune cell abundance recapitulated known immune phenotypes and suggested novel tissue-specific roles. Together, our findings provide a framework for understanding the molecular basis of variable expressivity in CDG and offer a resource for future functional and translational research.

### Statistics and reproducibility

The statistical methods utilized in each analysis are detailed within their corresponding sections. All statistical tests were performed using specialized R packages customized for each specific analysis.

## Data Availability

In this study, we utilized open-access data from healthy human tissues generated by the GTEx project (release v8). The open-access data, available through the GTEx Portal, include bulk gene expression values in TPM, haplotype-resolved expression counts from WASP-corrected RNA-seq alignments with phasing maintained across genes per sample, sample annotations, precomputed eQTLs, and snRNA-seq. We also used GTEx expanded subject phenotype data, accessed through the NCBI database of Genotypes and Phenotypes (dbGaP) under accession phs000424.v8.p2, in compliance with the NIH Genomic Data Sharing Policy.

## Code availability

The complete code required to replicate the analyses will be made available upon publication in a journal.

## Author Contributions

Cátia José Neves performed all analyses, generated the plots, and wrote the manuscript. Ana Rita Grosso designed the project. Ana Rita Grosso and Paula Videira supervised the study and critically revised the manuscript. Cátia José Neves and Rita Adubeiro Lourenço assigned CDG phenotypes to tissues, and António Gomes reviewed the phenotype assignments.

## Notes

https://gtexportal.org/home/

https://www.ensembl.org/index.html

https://www.ebi.ac.uk/gwas/

https://www.ncbi.nlm.nih.gov/clinvar/

https://www.malacards.org/

### Competing Interest Statement

The authors have declared no competing interest.

