## Supplementary material for "Tissue-Specific Regulatory and Expression Patterns of CDG-causative genes account to Phenotypic Variability": sup_figures

### Sup Figure 1

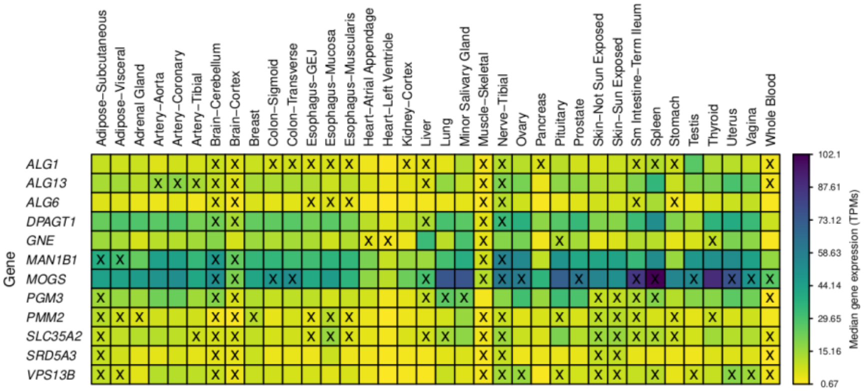


**Sup. Figure 1. Expression of CDG-causative genes across healthy tissues.** Heatmaps with the expression profiles of CDG-causative genes across various healthy tissues (median gene expression in Transcripts Per Million - TPM). The 'x' symbols highlight tissues most frequently affected by CDG types associated with these genes.

### Sup Figure 2

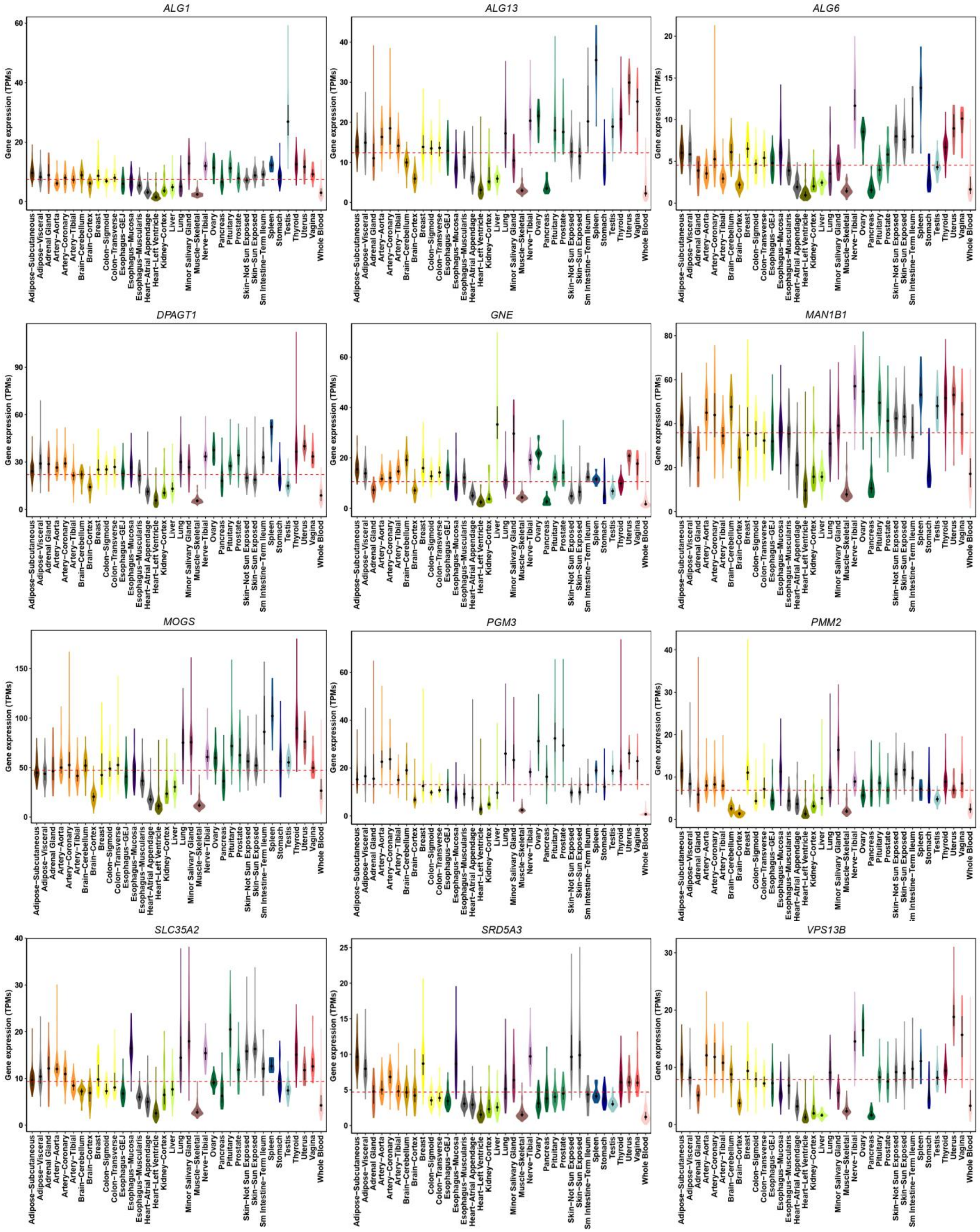


**Sup. Figure 2. Expression of CDG-causative genes across healthy tissues**. Distribution of gene expression levels (TPMs) for CDG-causative genes across healthy tissues. Each tissue group is represented by a unique color violin. The black dot represents the median and the thick black bar in the center represents the interquartile range. The horizontal red line represents the overall media across all tissues.

### Sup Figure 3

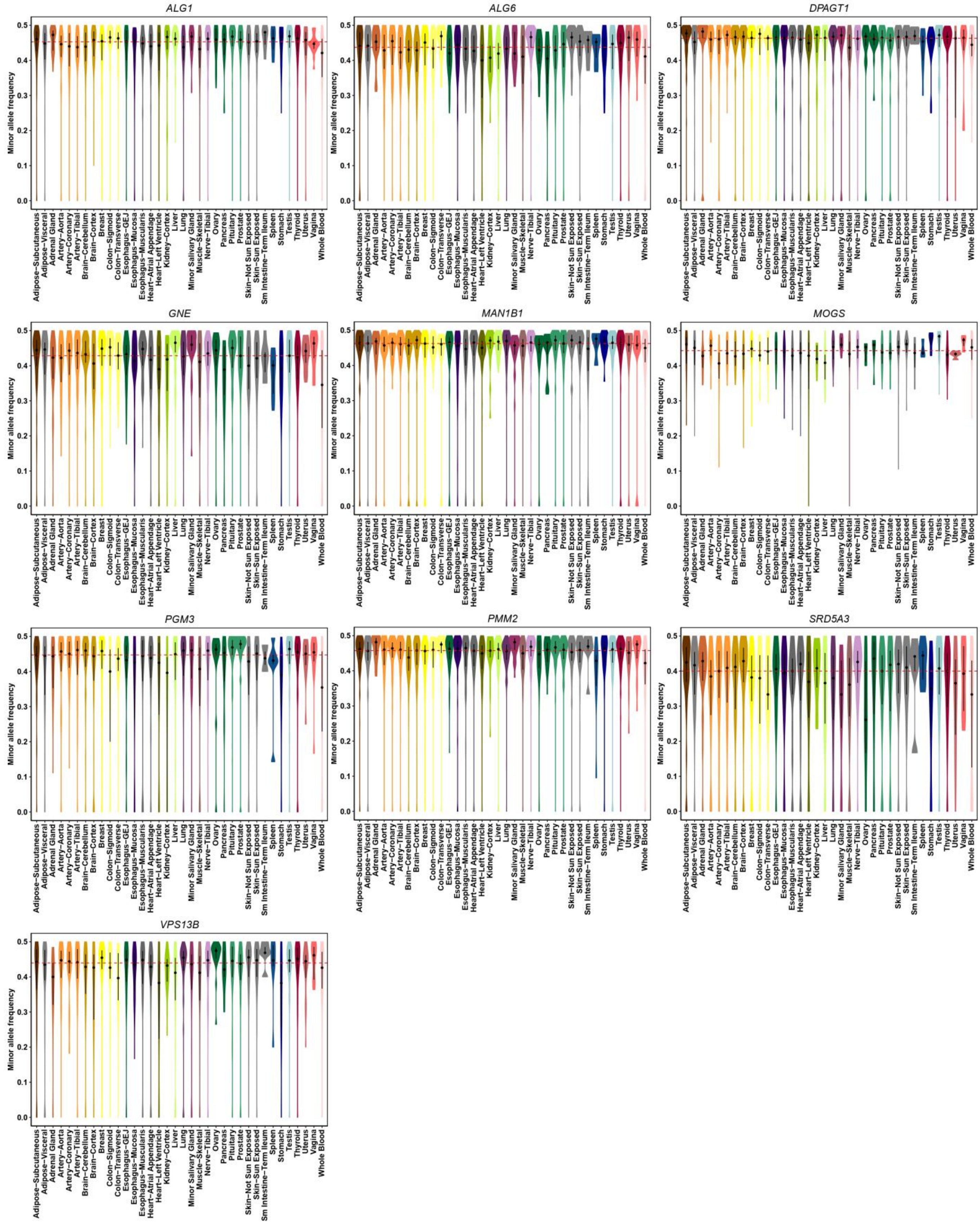


**Sup. Figure 3. Allelic expression of CDG-causative genes across healthy tissues.** Violin plots displaying the distribution and probability density of MAF for CDG-causative genes across various healthy tissues. Each tissue group is represented by a unique color violin. The black dot in the middle represents the median and the thick black bar in the center represents the interquartile range. The horizontal red line represents the overall median across all tissues.

### Sup Figure 4

## A

| Allele expression | Allele 1 ratios distribution of gene across at least 3 tissues (Kravitz et al., 2023) | Tissues with monoallelic expression (Alelle 1 ratio: [0-0.05] OR [0.95-1]) | Tissues with biallelic expression (Alelle 1 ratio: > 0.05 & < 0.95) | Alelle 1 ratio mean |
|---|---|---|---|---|
| **Biallelic balanced** | Binomial | 0 | >0 | >= 0.45 & <= 0.55 |
| **Biallelic biased** | Binomial | 0 | >0 | > 0.05 & < 0.45 OR > 0.55 & < 0.95 |
| **Tissue-specific monoallelic / Tissue-specific imprinting** | Binomial | >0 | >0 | _ |
| **Constitutive monoallelic/ Constitutive imprinting** | Binomial | >0 | 0 | _ |
| **Autosomal random monoallelic expression (aRME) / X-inactive** | β-Binomial | _ | _ | _ |

## B

| Positive control of allele expression type | Gene | Description |
|---|---|---|
| Biallelic balanced | MAPK1 | Subject GTEX-YFC4 with biallelic balanced (Kravitz et al. 2023) |
| | HNRNPA2B1<br>RAB11B<br>CSNK2B<br>RHOA | Housekeeping genes, with the most stable transcripts (Hounkpe et al. 2021) + hc-Biallelic (all tissues)(Kravitz et al. 2023) |
| Biallelic biased | ARL17A | Subject GTEX-YFC4 with biallelic biased (Kravitz et al. 2023) |
| Tissue-specific monoallelic/ Tissue-specific imprinting | LPAR6<br>UBE3A | Tissue-specific imprinting (Baran et al. 2015) |
| Constitutive monoallelic/ Constitutive imprinting | NAP1L5<br>SNRPN | Imprinting (Baran et al. 2015) |
| Autosomal random monoallelic expression | PARD6G | Subject GTEX-YFC4 with hc-RAE (Kravitz et al. 2023) |
| | GALNT2<br>PIGG<br>A4GALT | hc-RAE (all tissues)(Kravitz et al. 2023) + CDG gene |

## C

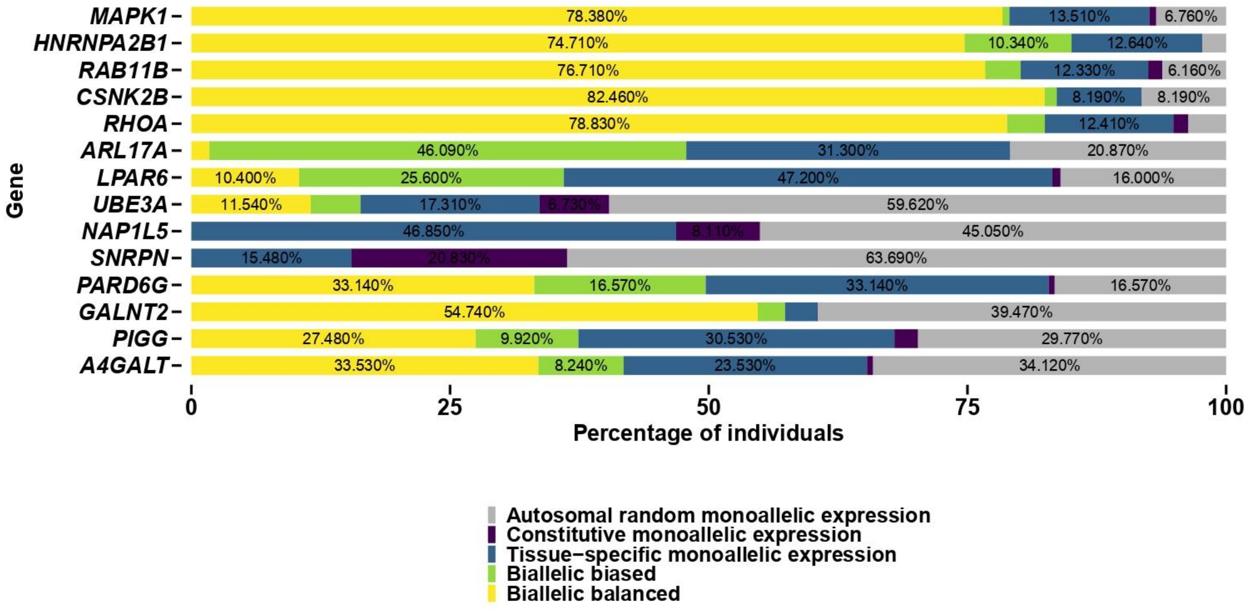


**Sup. Figure 4. Identifying various allelic expression patterns.** (A) Table displaying the parameters utilized to discern the allele expression type for the target gene, applying haplotype-phased RNA-seq data from the GTEx datasets across at least three tissues within a single individual. (B) Table with positive control genes to validate the parameters for differentiating allelic expression types. (C) Stacked barchart showing the percentage of individuals with different allelic expression types for each positive control gene.

### Sup Figure 5

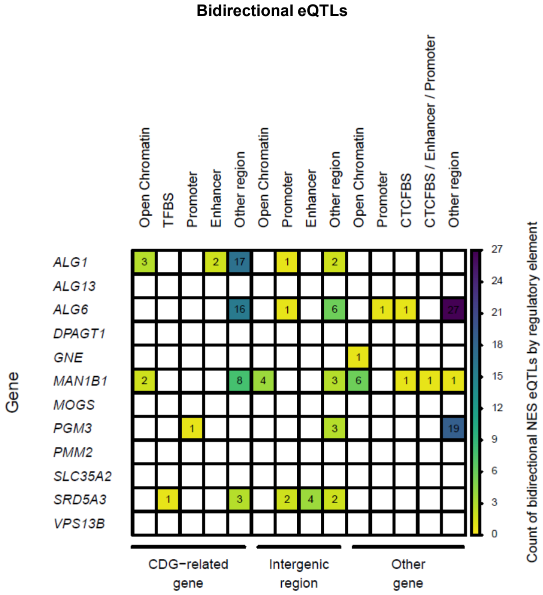


**Sup. Figure 5. Genomic Location of CDG-related eQTLs.** Heatmaps showing the number of eQTLs with bidirectional NES located in regulatory regions of the gene itself, in intergenic regions, or within another gene. Regulatory categories include open chromatin regions, transcription factor binding sites (TFBS), predicted promoters, predicted enhancers, CTCF binding sites (CTCFBS), and other regions.

### Sup Figure 6

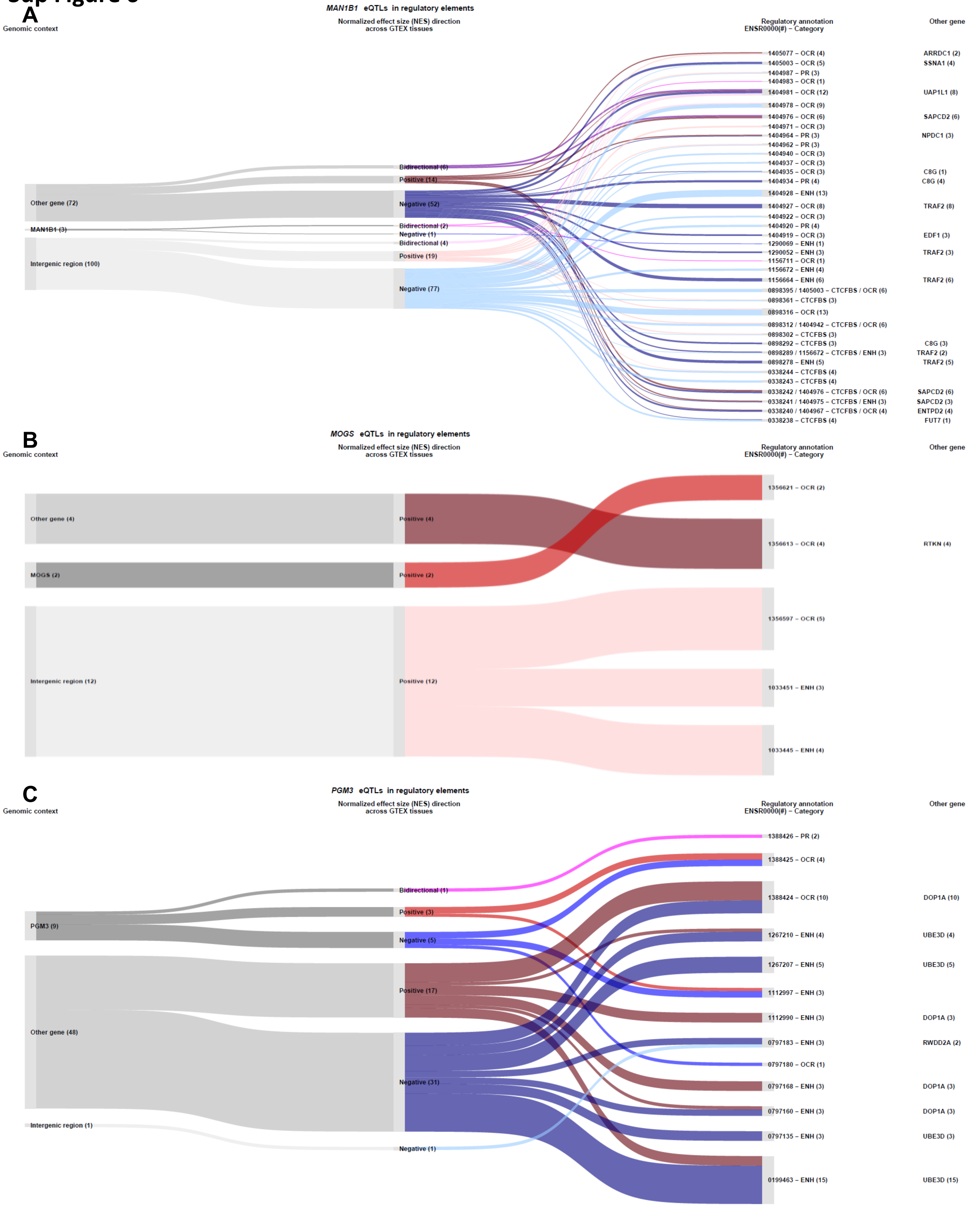


**Sup. Figure 6. Genomic Location of eQTLs associated with *MAN1B1*, *MOGS*, and *PGM3*.** (A-C) Sankey plots illustrating the number and distribution of eQTLs for the *MAN1B1* (A), *MOGS* (B) and *PGM3* (C) genes across regulatory regions. Regulatory categories include open chromatin regions (OCR), transcription factor binding sites (TFBS), predicted promoters (PP), predicted enhancers (ENH), CTCF binding sites (CTCFBS), and other regions. Displayed regulatory regions were selected based on their location and eQTL count: those overlapping the gene itself include ≥1 eQTL, while those overlapping another gene and/or an intergenic region include ≥3. Flows connect three eQTL classifications (nodes): (1) Genomic context - intergenic, within the gene itself, or another gene (grey shades); (2) NES direction across GTEx tissues - positive (red shades), negative (blue shades), or bidirectional (purple shades); (3) Regulatory annotation (Ensembl ID and category). Flow shading reflects genomic context: light for intergenic regions, medium for eQTLs within the gene itself, and dark for those within another gene. The final column lists the names of other genes, where applicable.

### Sup Figure 7

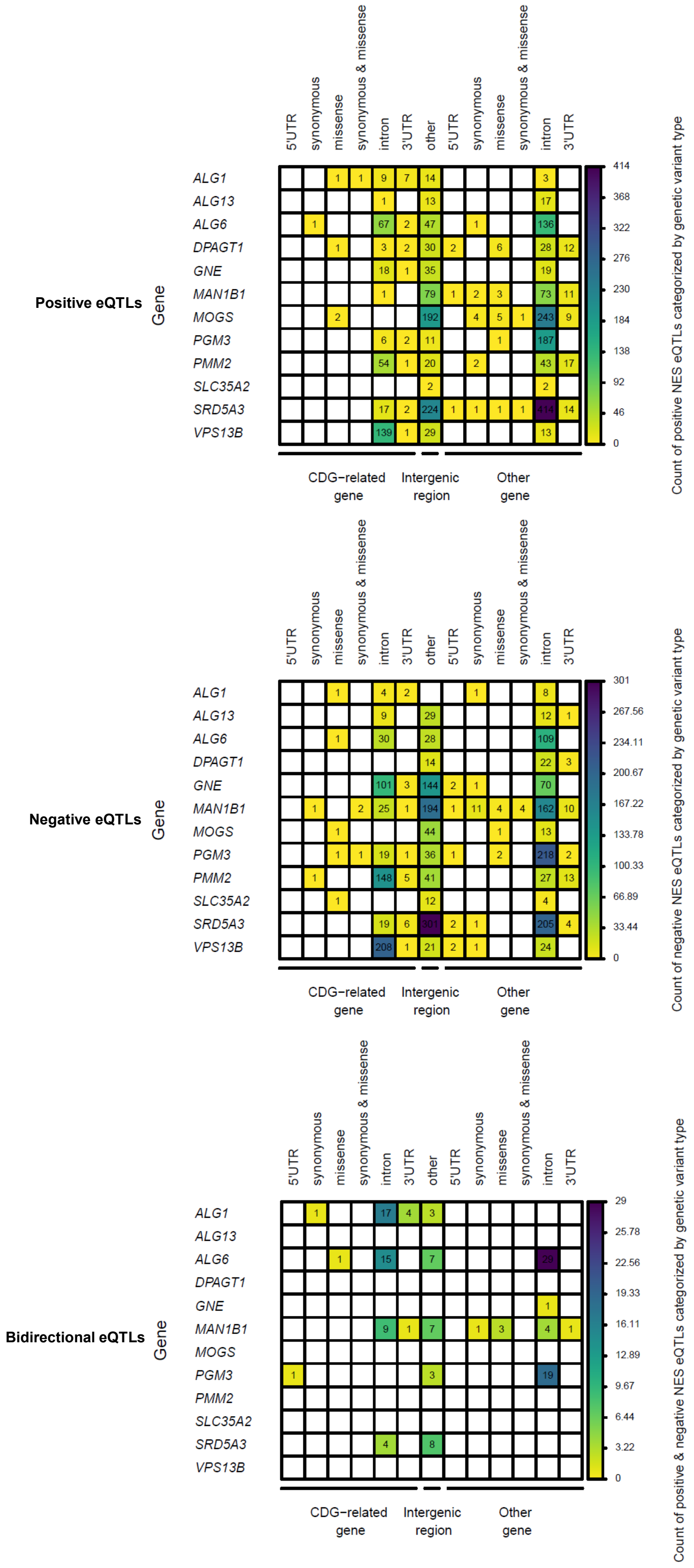


**Sup. Figure 7. Genetic variant types of CDG-related eQTLs.** Heatmaps showing the number of eQTLs with positive, negative, or bidirectional NES, grouped by genetic variant type. Variant types include 5′ untranslated region (5′UTR), synonymous, missense, combined synonymous and missense, intronic, 3′ untranslated region (3′UTR), and other variants. eQTLs are further categorized by genomic context: located within the gene itself, within another gene, or in an intergenic region.

### Sup Figure 8

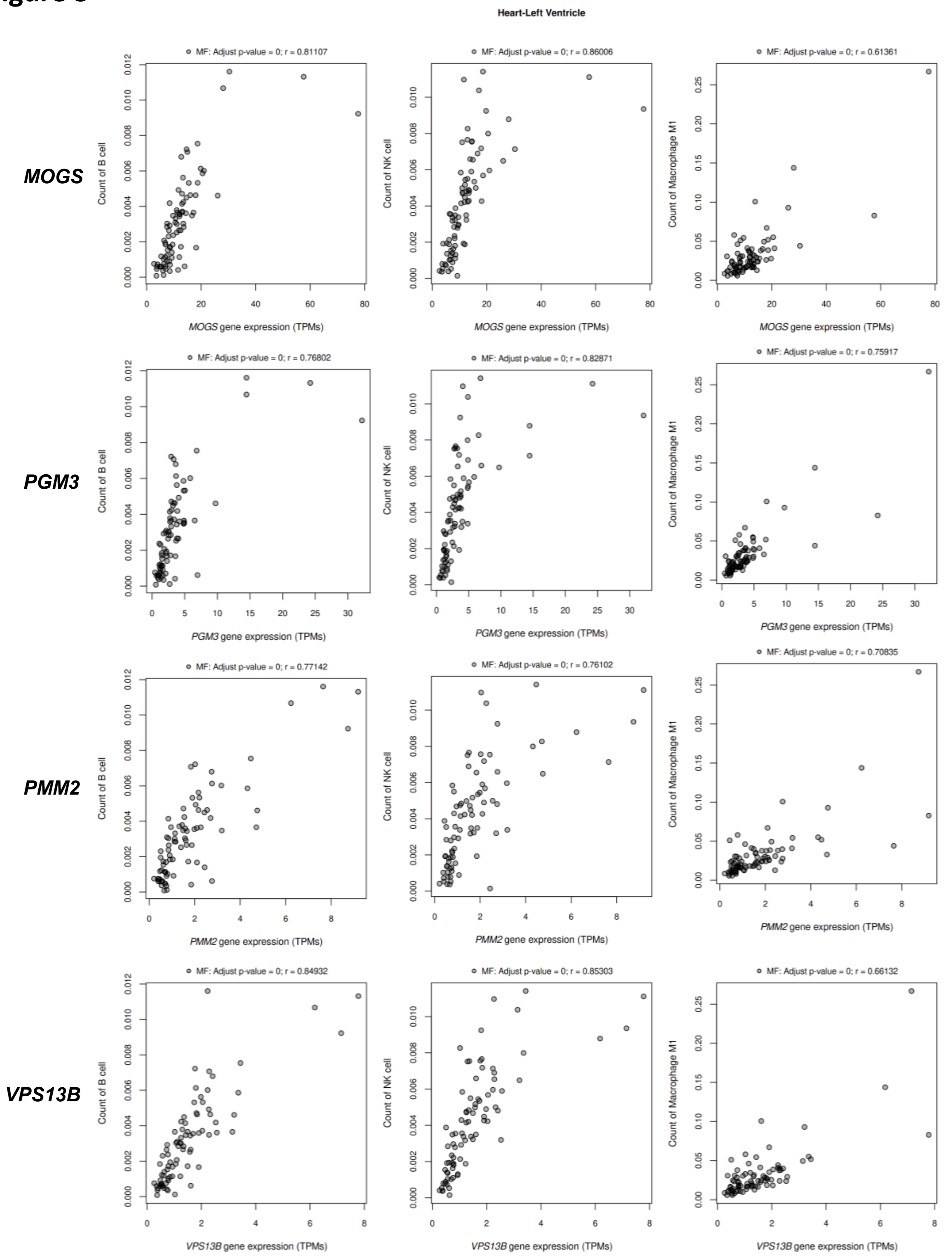


**Sup. Figure 8. Strong significant correlations between CDG-causative gene expression and immune cell abundance in healthy heart tissue.** Scatterplots showing strong positive correlations between the expression of *MOGS*, *PGM3*, *PMM2*, and *VPS13B* (TPM) and the abundance of B cells, NK cells, and M1 macrophages in healthy heart tissue. Correlations were considered significant if FDR-adjusted $p \leq 0.05$ and |Spearman r| ≥ 0.3.

### Sup Figure 9

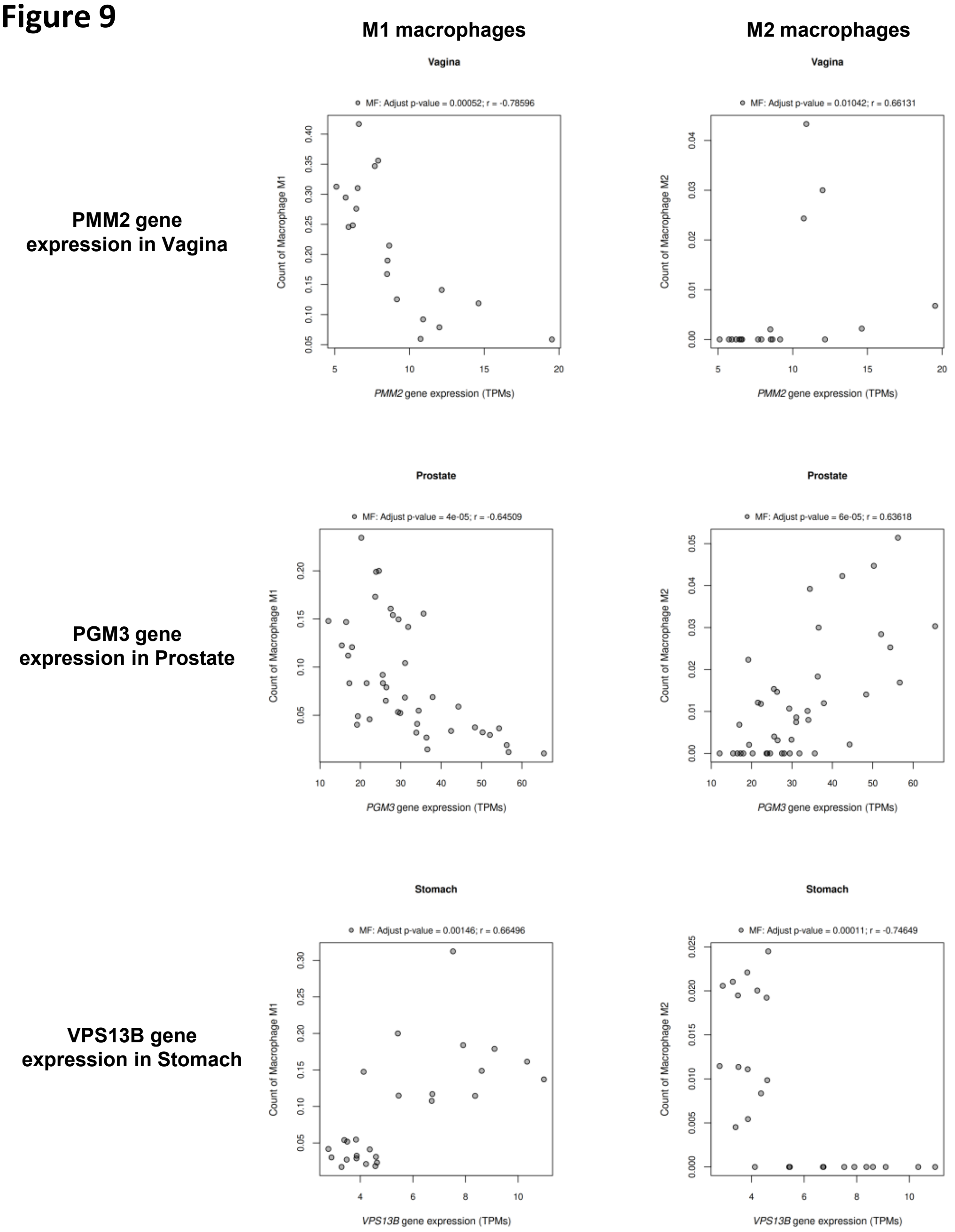


**Sup. Figure 9. Opposing correlations between M1 and M2 macrophage abundance and CDG-causative gene expression in healthy tissues.** Scatterplots showing inverse correlations between M1 and M2 macrophage abundance and the expression of *PMM2*, *PGM3*, and *VPS13B* (TPM) in healthy vagina, prostate, and stomach tissues, respectively. Correlations were considered significant if FDR-adjusted $p \leq 0.05$ and $|\text{Spearman } r| \geq 0.3$.

### Sup Figure 10

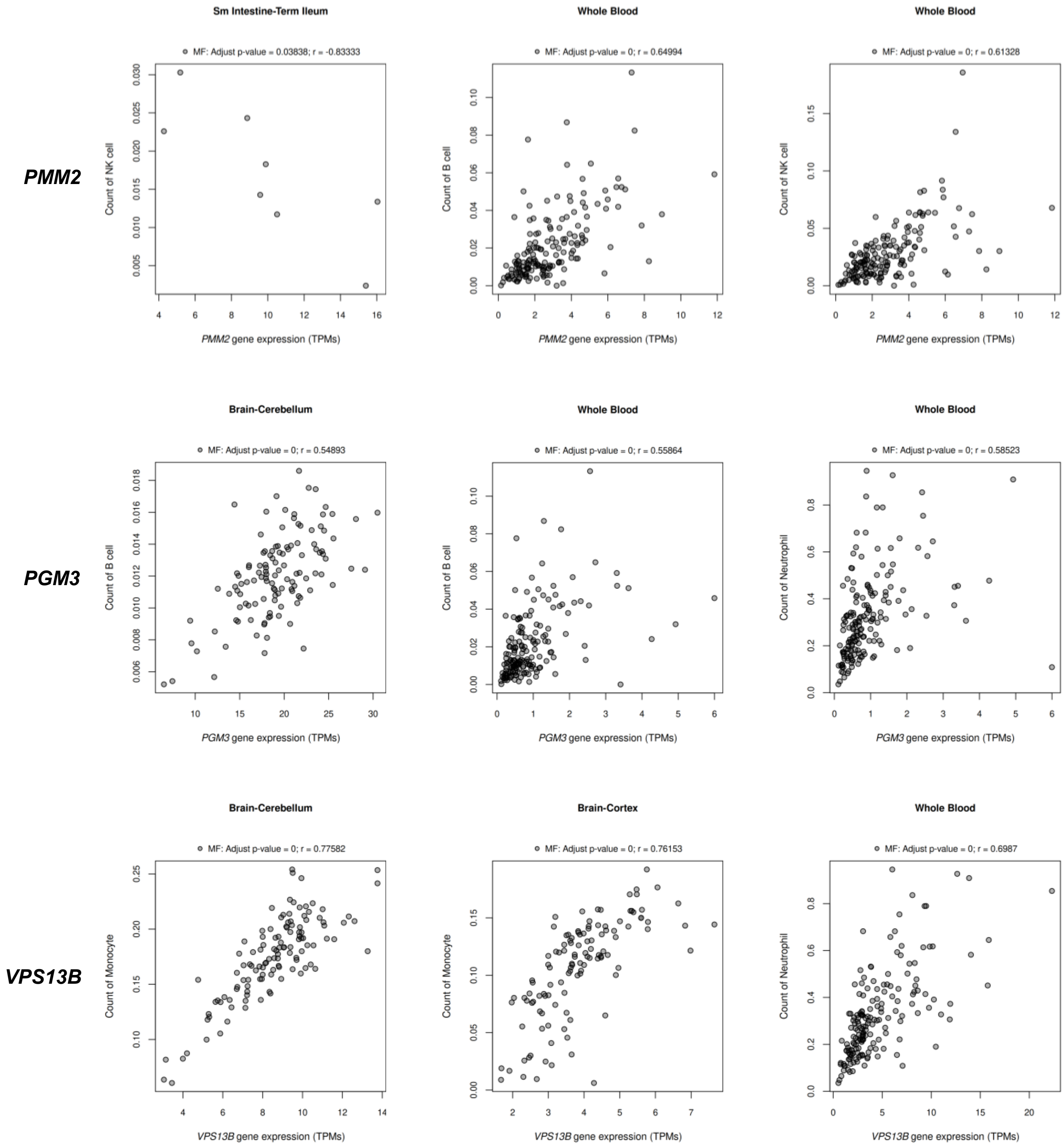


**Sup. Figure 10. Strong significant correlations between CDG-causative gene expression and immune cell abundance in healthy tissues frequently affected by the respective CDG type.** Scatterplots showing strong correlations between the expression of *PMM2*, *PGM3*, and *VPS13B* (TPM) and immune cell counts in healthy tissues commonly affected in their associated CDG types. Correlations were considered significant if FDR-adjusted $p \leq 0.05$ and |Spearman r| ≥ 0.3.

### Sup Figure 11

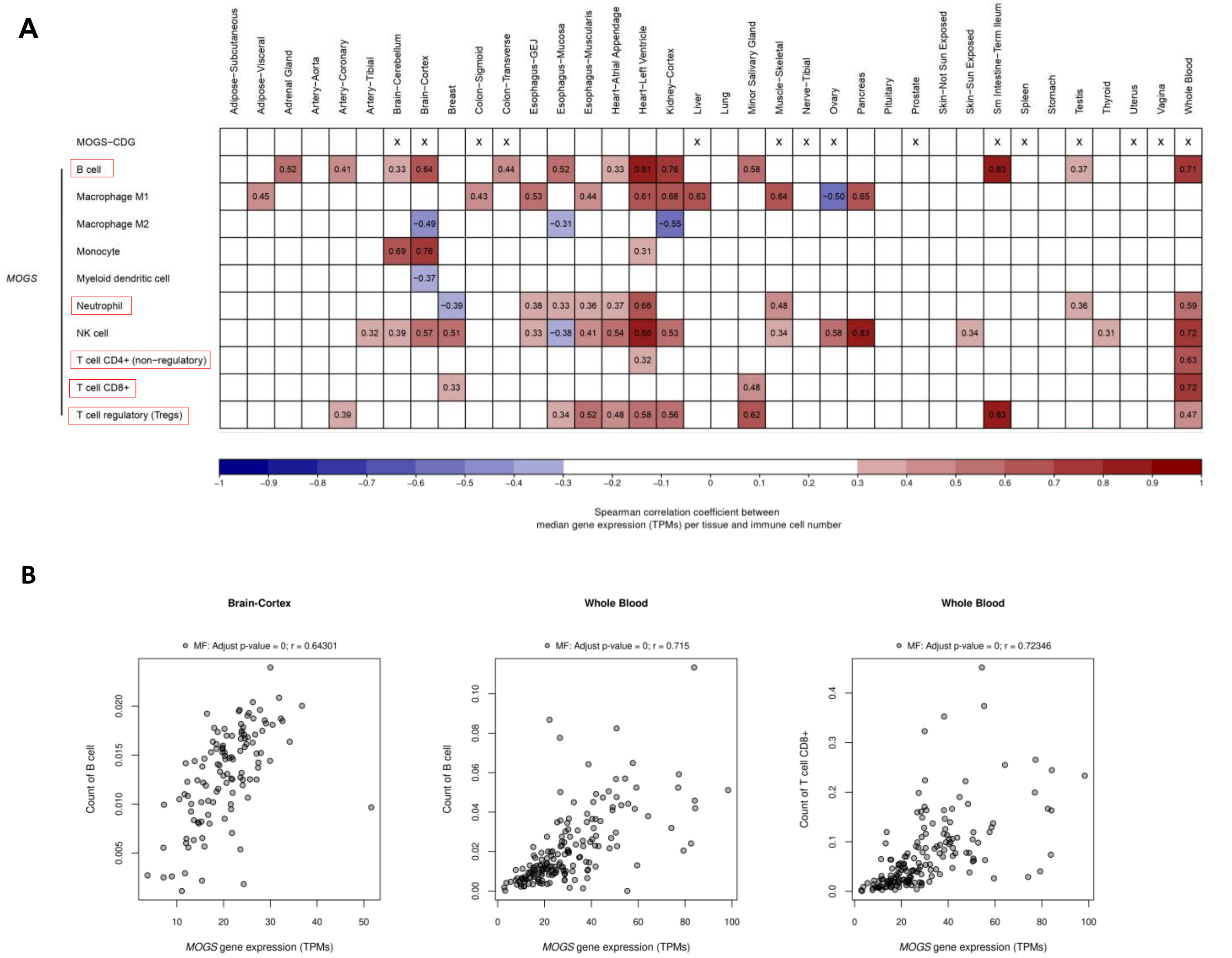


**Sup. Figure 11. Immune cell types associated with the expression of *MOGS*.** (A) Heatmap showing significant correlations between median *MOGS* gene expression (TPM) and immune cell counts across tissues (FDR-adjusted $p \leq 0.05$ and |Spearman r| ≥ 0.3). Correlation strength is indicated by color intensity for positive (red) and negative correlation (blue). White indicates no significant correlation. An 'x' in the top row marks tissues frequently affected in MOGS-CDG. (B) Scatterplots showing strong correlations between *MOGS* expression (TPM) and immune cell counts in healthy tissues commonly affected in MOGS-CDG. Correlations were considered significant if FDR-adjusted $p \leq 0.05$ and |Spearman r| ≥ 0.3.

### Sup Figure 12

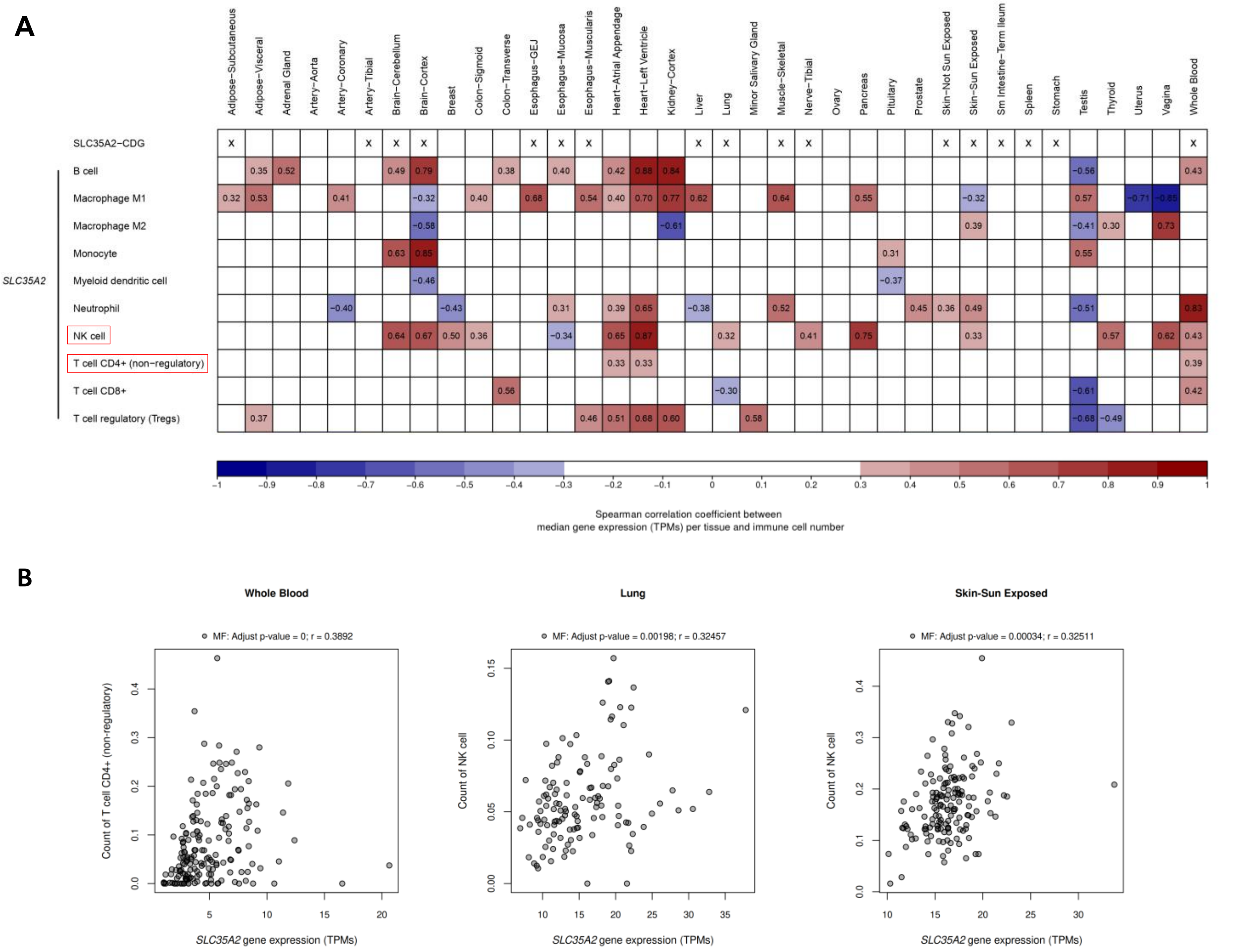


**Sup. Figure 12. Immune cell types associated with the expression of *SLC35A2*.** (A) Heatmap showing significant correlations between median *SLC35A2* gene expression (TPM) and immune cell counts across tissues (FDR-adjusted $p \leq 0.05$ and |Spearman r| ≥ 0.3). Correlation strength is indicated by color intensity for positive (red) and negative correlation (blue). White indicates no significant correlation. An 'x' in the top row marks tissues frequently affected in *SLC35A2*-CDG. (B) Scatterplots showing strong correlations between *SLC35A2* expression (TPM) and immune cell counts in healthy tissues commonly affected in *SLC35A2*-CDG. Correlations were considered significant if FDR-adjusted $p \leq 0.05$ and |Spearman r| ≥ 0.3.

### Sup Figure 13

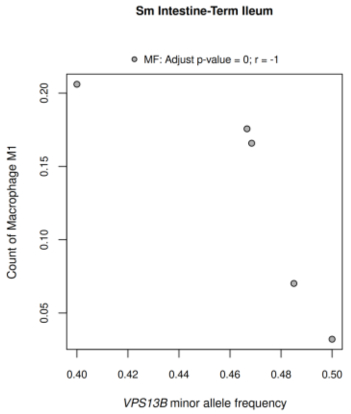


**Sup. Figure 13. Immune cell types associated with the minor allele frequency of *VPS13B*.** Scatterplot showing a strong negative correlation between *VPS13B* minor allele frequency and M1 macrophages in healthy terminal ileum. Correlations were considered significant if FDR-adjusted *p* ≤ 0.05 and |Spearman r| ≥ 0.3.
